# Dynamic network-guided CRISPRi screen reveals CTCF loop-constrained nonlinear enhancer-gene regulatory activity in cell state transitions

**DOI:** 10.1101/2023.03.07.531569

**Authors:** Renhe Luo, Jielin Yan, Jin Woo Oh, Wang Xi, Dustin Shigaki, Wilfred Wong, Hyunwoo Cho, Dylan Murphy, Ronald Cutler, Bess P. Rosen, Julian Pulecio, Dapeng Yang, Rachel Glenn, Tingxu Chen, Qing V. Li, Thomas Vierbuchen, Simone Sidoli, Effie Apostolou, Danwei Huangfu, Michael A. Beer

## Abstract

Comprehensive enhancer discovery is challenging because most enhancers, especially those affected in complex diseases, have weak effects on gene expression. Our network modeling revealed that nonlinear enhancer-gene regulation during cell state transitions can be leveraged to improve the sensitivity of enhancer discovery. Utilizing hESC definitive endoderm differentiation as a dynamic transition system, we conducted a mid-transition CRISPRi-based enhancer screen. The screen discovered a comprehensive set of enhancers (4 to 9 per locus) for each of the core endoderm lineage-specifying transcription factors, and many enhancers had strong effects mid-transition but weak effects post-transition. Through integrating enhancer activity measurements and three-dimensional enhancer-promoter interaction information, we were able to develop a CTCF loop-constrained Interaction Activity (CIA) model that can better predict functional enhancers compared to models that rely on Hi-C-based enhancer-promoter contact frequency. Our study provides generalizable strategies for sensitive and more comprehensive enhancer discovery in both normal and pathological cell state transitions.

## Main text

Many consortia have made important progress in mapping putative enhancers based on chromatin accessibility and protein binding in a wide range of cell types and tissues ^1^. Harnessing the atlas of enhancers predicted from chromatin features, functional interrogation with large-scale CRISPR screens has successfully identified some enhancers with relatively strong impacts on gene expression in various cell lines ^2–15^. However, comprehensive enhancer discovery remains challenging. In some cases, enhancer perturbation only causes temporary phenotypes ^7, 8^. In other cases, the effect of enhancer perturbation is mitigated by the activity of “shadow” or redundant enhancers ^15–21^. Furthermore, most genetic variants associated with common human diseases show relatively modest effects on expression in reporter assays ^22, 23^, posing a challenge for identifying causal disease variants. A critical missing element is a quantitative model for how multiple enhancers work together at each locus to respond to physiological stimuli and experimental perturbations in a nonlinear way through altering gene regulatory network (GRN) activity. GRNs control cell states through a handful of core transcription factors (TFs), which both self-regulate and cooperatively regulate each other through core enhancers ^24^. Therefore, we focused on deconstructing the core enhancers’ activity in the GRN through determining the impact of their perturbation on the cell state. We developed a quantitative GRN model to simulate the dynamic process of core circuit establishment. The results predict that cell states are more susceptible to core enhancer perturbations during dynamic cell state transitions compared to post-transition when the GRN has been fully established. These results prompted us to use human embryonic stem cell (hESC) guided differentiation as a platform to perturb enhancers during cell state transitions. We designed a dCas9-KRAB-based CRISPRi screen to interrogate 394 putative enhancers surrounding 10 core TF loci spanning a total of 40 Mb genomic regions during hESC to definitive endoderm (DE) differentiation. Using *SOX17* as the DE cell state readout ^25^, we identified multiple enhancers (4 to 9 per locus) for each of the core DE TFs (*EOMES*, *GATA6*, *MIXL1*, and *SOX17*). This affirms the feasibility of using a single core gene as the readout for discovering functional core enhancers during cell state transitions. The sensitive screening strategy also uncovered 12 enhancers >100 kb away from the transcription start site (TSS) of the target gene, demonstrating the need for enhancer discovery beyond the immediate linear neighborhood. The relatively comprehensive discovery of functional enhancers allowed us to develop a CTCF loop-constrained Interaction Activity (CIA) model that outperformed previous Hi-C contact-based enhancer prediction methods ^4, 26–28^. Our network-guided core enhancer mapping strategy during cell state transitions and the CIA enhancer prediction model provide a framework for systematic enhancer discovery applicable not only to normal development but also to pathological conditions such as diabetes and cancer.

## Dynamic GRN model predicts temporal sensitivity to enhancer perturbation

We sought to develop a dynamic GRN model to study the temporal and threshold-dependent requirements for enhancers during cell state transitions. Our previous studies of cell state transitions ^24, 25, 29, 30^ and sequence-based modeling ^24, 31, 32^ of a broad range of ENCODE epigenomic profiling data have identified key features of this model. Machine learning applied to chromatin accessible peaks identifies a small set of 5-10 lineage determining core TFs in each cell type whose binding sites can predict chromatin accessible peaks to a high degree of accuracy ^24, 31, 32^. Each chromatin accessible peak contains combinations of multiple binding sites for these core TFs. Thus, the lineage determining core TFs cooperatively auto-regulate each other through multiple enhancers flanking each core TF gene and they co-regulate downstream peripheral genes (Fig.1a; Extended Data Fig.1a). This results in highly nonlinear regulation of gene expression. Since these core TFs determine the network state, we will describe the activity of the network with a time-dependent state variable, *ψ*(*t*), whose amplitude reflects the activity of the core TFs. We use ‘network state’ and ‘cell state’ essentially interchangeably. The key features of this model are as follows: each gene is expressed at an activated (*e*_o_) or basal level (*e*_"_), depending on the activity of its flanking enhancers. Each gene will be in the activated state with probability *p*_on_ ∼ *cf*(*ψ*) and in the basal state with *p*_off_ ∼ *b*, where *f*(*ψ*) is a nonlinear function of the core TF activity and reflects cooperativity at the enhancers, and whose activity can be modulated through the parameter *c*, (e.g., via CRISPRi). In addition, we allow a time-dependent differentiation stimulus *δ*(*t*) that acts at the enhancers through a separate mechanism (e.g., differentiation signaling) to stimulate the transition, so *p*_on_ ∼ *cf*(*ψ*) + *δ*(*t*). Finally, we add degradation or export of the TFs (−*rψ*) and a stochastic noise term, *ξ*(*t*), so the final network state equation is:

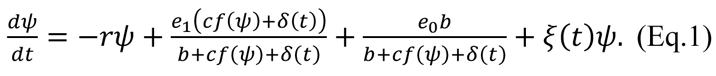

**Figure 1.**
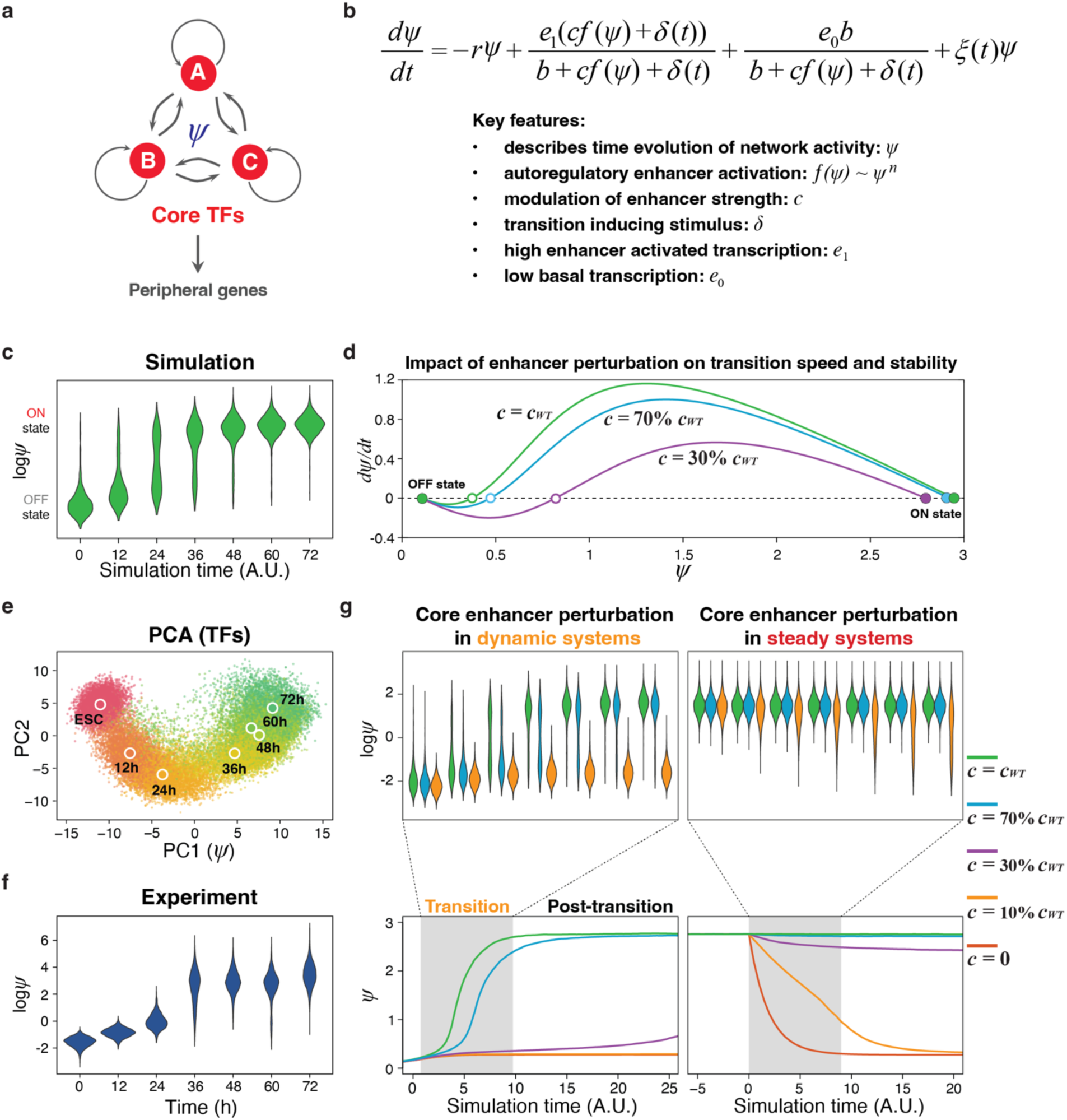
Dynamic Gene Regulatory Network model. **a**, The schematic of core circuit in the GRN. The core TFs cooperatively auto-regulate each other and co-regulate downstream peripheral genes. **b**, The equation of the dynamic gene regulatory network model. **c**, The violin plots of core circuit establishment during cell state transition by simulation. **d**, Plot of *dψ*⁄*dt* vs *ψ* showing cell state transition with different enhancer strengths. The green line represents total enhancer strength without perturbation. The cyan line represents total enhancer strength reduced to 70% of full strength by perturbation. The purple line represents total enhancer strength reduced to 30% of full strength by perturbation. **e**, PCA analysis of all TFs from scRNA-seq data during hESC-DE transition. Each dot represents a cell collected every 12h during hESC-DE transition. The large dots represent the average of all cells from the same time point. **f**, The violin plots of core circuit establishment during hESC-DE transition by scRNA-seq experiments. **g**, The comparison between the same enhancer perturbation strength during cell state transition or at steady state. The line plots represent the median of simulation results. The violin plots correspond to a zoomed in time interval denoted by the grey box on the line plots. The green, cyan, purple, yellow and orange lines represent 100%, 70%, 30%, 10% and 0% of the original total enhancer strength respectively.

The simplifications that lead to this one-dimensional network state equation, where *ψ*(*t*) is a scalar representing the degree to which the network is activated, are described in Methods. In the strongly cooperative limit, the enhancers together have an activity *f*(*ψ*) ≈ *ψ*^$^ where *B* is a typical number of TFs binding at each enhancer. For concreteness and simplicity, we will take *B* = 3, but our conclusions are robust for *n* ≥ 3. Stochastic simulation of this model (Methods) produces distributions of cells with either high or low network activity (Fig.1c). This can be understood from stability analysis of this model, by plotting *dψ*⁄*dt* vs *ψ* (Fig.1d). Over a wide parameter range, the network has three fixed points where *dψ*⁄*dt* = 0. Two of these are stable states: the OFF state, with low activity where basal activation balances degradation, and the ON state, with high activity where enhancer-driven transcriptional activation balances degradation. There is also an intermediate unstable fixed point (Fig.1d), above which the network activity will transition to the ON state, otherwise, it will fall to the OFF state. The simulation results are in good agreement with single-cell RNA sequencing (scRNA-seq) experiments sampling every 12 hours (12h) during hESC-DE differentiation (Fig.1c, e, f). To generalize the definition used in the equation, we used the projection of the expression of all TFs along principal component analysis (PCA) component 1 to measure the network state in each cell (Fig.1e; Extended Data Fig.1b, c). The scRNA-seq results show a bi-stable distribution of cell states during differentiation: a pre-transition steady state, from 0h to 24h, and a post-transition steady state 48h to 72h, when transcriptome profiles are relatively similar over time, and a transitional period, from 24h to 48h, when transcriptome profiles are changing more rapidly (Fig.1e, f; Extended Data Fig.1d). This transition behavior is also predicted in the simulation results (Fig.1c, d). The consistency between simulation and experimental results suggests that the simple dynamic network model is nevertheless capturing the key features of the establishment of the core circuit in a GRN.

To quantitatively model enhancer perturbation during the cell state transition, we decreased the total enhancer strength (*c*) to varying degrees (with the same stimulus strength *δ*(*t*)) and compared the simulation results. Mildly decreasing the total enhancer strength, which mimics perturbing one of the multiple core enhancers flanking a core TF, has a very weak effect on both the ON and OFF steady states, but has a much stronger effect on the network activity required to transition between states (the unstable fixed point where *dψ*⁄*dt* = 0 in Fig.1d), indicating mild enhancer perturbation could delay the cell state transition without dramatically changing the final network state (Fig.1d, g left). We also mimicked the enhancer perturbation in a steady state condition by decreasing the total enhancer strength (*c*) after cells have reached the steady state. Compared with the simulation during the forward cell state transition, the same enhancer perturbations show much weaker effects in steady state (Fig.1g). These results indicate that an optimal time window exists during cell state transitions, which could be utilized to increase the sensitivity of enhancer perturbation screens over screens conducted at a steady state ^2–10^. We further validated the results of the simple GRN network model (Eq.1) with stochastic Gillespie simulations ^33, 34^ (formally valid in the limit of small numbers of TFs) under the same nonlinear auto-regulatory assumptions. These simulations reproduce the main findings of the sensitivity to enhancer perturbation during the transition, temporal delay in the transition, and insensitivity to enhancer perturbation after the GRN is fully activated in the ON state (Extended Data Fig.1e; Methods).

## A dynamic network-guided CRISPRi screen identified core enhancers during cell state transition

We applied our modeling results to discover core enhancers in cell state transitions using hESC-DE differentiation as a test case, for which core enhancers remain incompletely defined ^35^. Optimization of our existing differentiation protocol ^25^ allowed us to reproducibly generate >95% SOX17^+^/CXCR4^+^ DE cells within 72 hours after the initiation of DE differentiation (DE-72h) (Extended Data Fig.2a; Methods). To identify core enhancers, we first took a systematic approach to define core TFs (Fig.2a). Since *OCT4* (*POU5F1*), *NANOG* and *SOX2* are well-known TFs essential for the ESC identity ^36^, we focused on identifying core TFs for the hESC-DE transition. We analyzed the genes required for hESC-DE transition identified from our unbiased genome-scale CRISPR-Cas9 screening data ^25^ (Fig.2b), and selected four core DE TFs (*EOMES*, *MIXL1*, *GATA6*, *SOX17*) and three signaling TFs (*SMAD2*, *SMAD4*, and *JUN*) (Fig.2c; Extended Data Fig.2b). The DE and ESC TFs showed opposing changes in gene expression during hESC-DE transition with corresponding changes in their regulatory activities predicted by gkm-SVM ^24, 31^ trained on the ATAC-seq data (Extended Data Fig.2c-f). Analysis of gene expression from scRNA-seq data (collected every 12h during hESC-DE transition) using the Pearson correlation as the distance metric for UMAP visualization further demonstrated that the expression patterns of DE and ESC TFs clustered separately, and they each correlated amongst themselves (Fig.2c; Extended Data Fig.2f; Methods). Compared to the ESC and DE TFs, the signaling TFs showed less dynamic changes in transcriptional and regulatory activities during differentiation (Extended Data Fig.2c-f). We further examined biochemical cooperation between the core DE TFs through chromatin immunoprecipitation followed by mass spectrometry (ChIP-MS) for EOMES, GATA6 and SOX17. Our analysis revealed that all three TFs interacted with each other, and they also interacted with many common partners including the DE TF MIXL1 and the signaling TF SMAD4 (Fig.2d; Extended Data Fig.2g; Extended Data table 1). In summary, we identified 10 core TFs for the study, starting with functional genomics data and corroborating with gene expression, chromatin accessibility and proteomics findings (Fig.2e).

**Figure 2.**
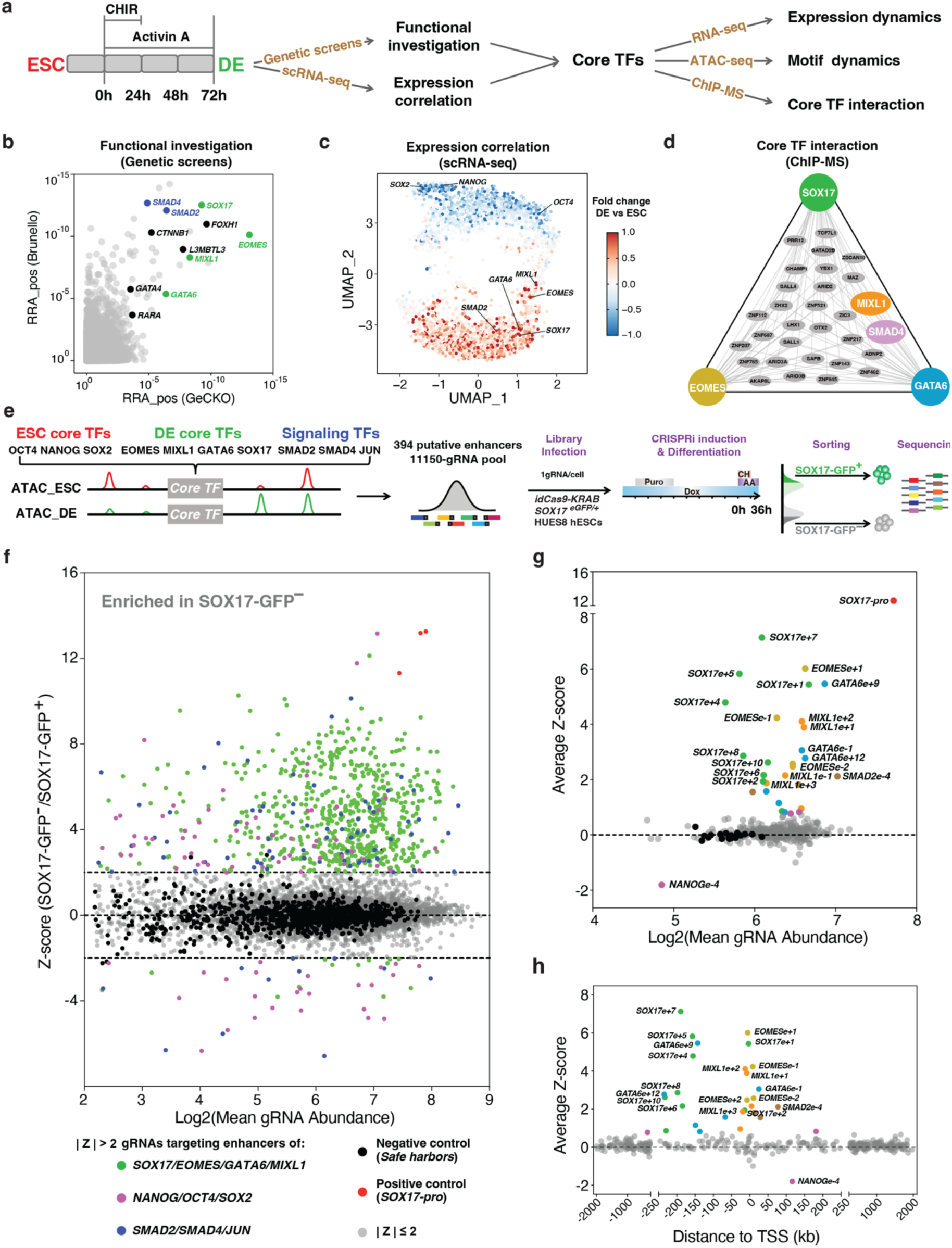
A dynamic network-guided enhancer screen identified core enhancers during hESC-DE cell state transition. **a**, The schematic of using a systematic approach to identify the core TFs during hESC-DE transition. **b**, MAGeCK robust ranking aggregation (RRA) scores for positive hits in two genome-scale DE screens from Li et al ^25^. We overlapped the top 100 hits from each screen, there were 11 TFs (labeled). DE core TFs and signaling TFs identified for the network-guided core enhancer perturbation screen are shown in green and blue, respectively. **c**, The expression correlation analysis from scRNA-seq of hESC-DE transition. Each dot represents a significant expressed gene during the hESC-DE transition (Methods). The distance between each pair of genes is defined by Pearson correlation. The color scale is defined from the expression fold change from ESS to DE-72h. **d**, The common protein interactive partners between core TFs identified by ChIP-MS using EOMES, GATA6 and SOX17 as baits during hESC-DE transition. **e**, The design of dynamic network-guided core enhancer perturbation screening. **f**, The scatter plot of gRNA Z-score distribution from the screen. Each dot represents an individual gRNA. Dashed lines indicate the threshold of |Z-score| = 2. Gray dots represent gRNAs of |Z-score| ≤ 2. Black dots represent negative control gRNAs. Red dots represent positive control gRNAs. Green dots represent gRNAs of |Z-score| > 2 targeting on *SOX17*, *EOMES*, *GATA6* and *MIXL1* loci. Purple dots represent gRNAs of |Z-score| > 2 targeting on *NANOG*, *OCT4* and *SOX2* loci. Blue dots represent gRNAs of |Z-score| > 2 targeting on *SMAD2*, *SMAD4* and *JUN* loci. **g**, The scatter plot of average Z-score distribution of each putative enhancer region from the screening. Each dot represents an individual region. Black dots represent negative controls. 29 enhancer hits are labeled by different colors representing the core TFs they are surrounding. **h**, Scatter plots showing the distance of 29 enhancers to the TSS of their target genes.

To discover core enhancers, we first examined 4 Mb genomic regions surrounding each of the 10 core TFs, and identified 394 regions that are accessible at either ESC or DE stage based on ATAC-seq (excluding the promoter regions). We then designed a tiling gRNA lentiviral library to target these regions with a total of 11,050 gRNAs (including 1,100 negative controls targeting safe harbor loci ^37^ and 3 positive controls targeting the *SOX17* promoter, Fig.2e; Extended Data Fig.4a-d; Extended Data table 2). We generated an hESC line with a doxycycline-inducible dCas9-KRAB cassette and a DE lineage reporter *SOX17^eGFP/+^* (Extended Data Fig.3a-d; Methods) and infected the cells with the gRNA library at a multiplicity of infection (MOI) of ∼0.3 to ensure that most infected cells received a single gRNA (Fig.2e). Based on flow cytometric analysis for SOX17-GFP expression during differentiation (Extended Data Fig.2a), as well as PCA analysis of scRNA-seq, RNA-seq and ATAC-seq data, we identified DE-36h as an optimal mid-transition point for interrogation of enhancer perturbation effects (Fig.1e, f; Extended Data Fig.2c-f). SOX17-GFP^+^ (top 20%) and SOX17-GFP^−^ (bottom 20%) cells were isolated through fluorescence-activated cell sorting (FACS) for gRNA enrichment analysis (Fig.2e; Methods). We calculated the Z-score for each gRNA based on its logarithm of fold change (log2FC) in the SOX17-GFP^−^ versus the SOX17-GFP^+^ cells (Fig.2f). Most gRNAs in the same hit regions had similar Z-scores (Extended Data Fig.4e, f), supporting that the screen is both sensitive and robust. Through calculating the average gRNA Z-score for each region, we discovered 29 enhancer hits with Z-scores ranging from 0.75 to 7.14 (Fig.2g; Extended Data table 3). Relatively few enhancers were found for the ESC and signaling TFs, likely reflecting that the ESC TFs mainly exert their regulatory effects at the ESC stages and that the signaling TFs are primarily regulated post-transcriptionally (e.g., through protein phosphorylation). In contrast, we discovered many enhancers for the core DE TFs, ranging from 4 to 9 for each gene (Fig.2g; Extended Data Fig.4f; Extended Data table 3). These findings demonstrate the high sensitivity of the screening strategy utilizing a dynamic transition and a single gene readout (e.g., *SOX17* for the DE identity) for discovery of core enhancers in multiple loci. Most of these enhancers were previously unknown, and their discovery expands the atlas of regulatory elements required for human development and provides a basis for understanding the complex gene regulatory networks that govern cell state transitions. Furthermore, 41% of the identified core enhancers are more than 100 kb away from the TSS of the cognate gene (Fig.2h), highlighting the need for examining putative enhancers in relatively broad genomic windows.

## Validation of core enhancers using CRISPRi perturbation and CRISPR-Cas9 mediated deletion

We selected 20 top enhancer hits for validation. Individual CRISPRi perturbations of all 20 hits resulted in significantly reduced numbers of SOX17-GFP^+^ cells at DE-36h based on flow cytometric analysis (Fig.3a, b). In addition to the cell state readout, we also confirmed the down-regulation of the corresponding cognate gene expression after perturbation by RT-qPCR analyses (Extended Data Fig.5a). In sharp contrast to the strong phenotypes observed at DE-36h, at DE-72h, the perturbations of most of the top 20 enhancers had little or no impact on the induction of SOX17-GFP^+^/CXCR4^+^ cells or their cognate gene expression (Fig.3a-d; Extended Data Fig.5a, b), which is reminiscent of the temporarily phenotypic enhancers previously described ^7, 8^. To investigate the enhancer perturbation effect on hESC-DE transition with a finer temporal resolution, we focused on two *GATA6* enhancers (*GATA6e+9* and *GATA6e+12*) and measured the differentiation efficiency based on SOX17 expression every 6 hours. Consistent with our dynamic GRN model (Fig.1g), the results show an exquisite temporal sensitivity to enhancer perturbations: a significant impact was observed only in a narrow time window (around DE-36h) during the transition (Fig.3e, f). In summary, these validation results show that the expression of core TF genes is frequently regulated by multiple enhancers, ensuring robustness in the regulatory network. Perturbing a single enhancer can significantly decrease target gene expression and delay cell-state transitions, but most perturbations do not substantially alter the post-transition state. The consistency of our experimental results with our dynamic GRN model suggests a cooperative autoregulatory mechanism underlying this effect. Together they support the utility of cell state transitions in perturbation screens as a generalizable approach for sensitized enhancer discovery.

**Figure 3.**
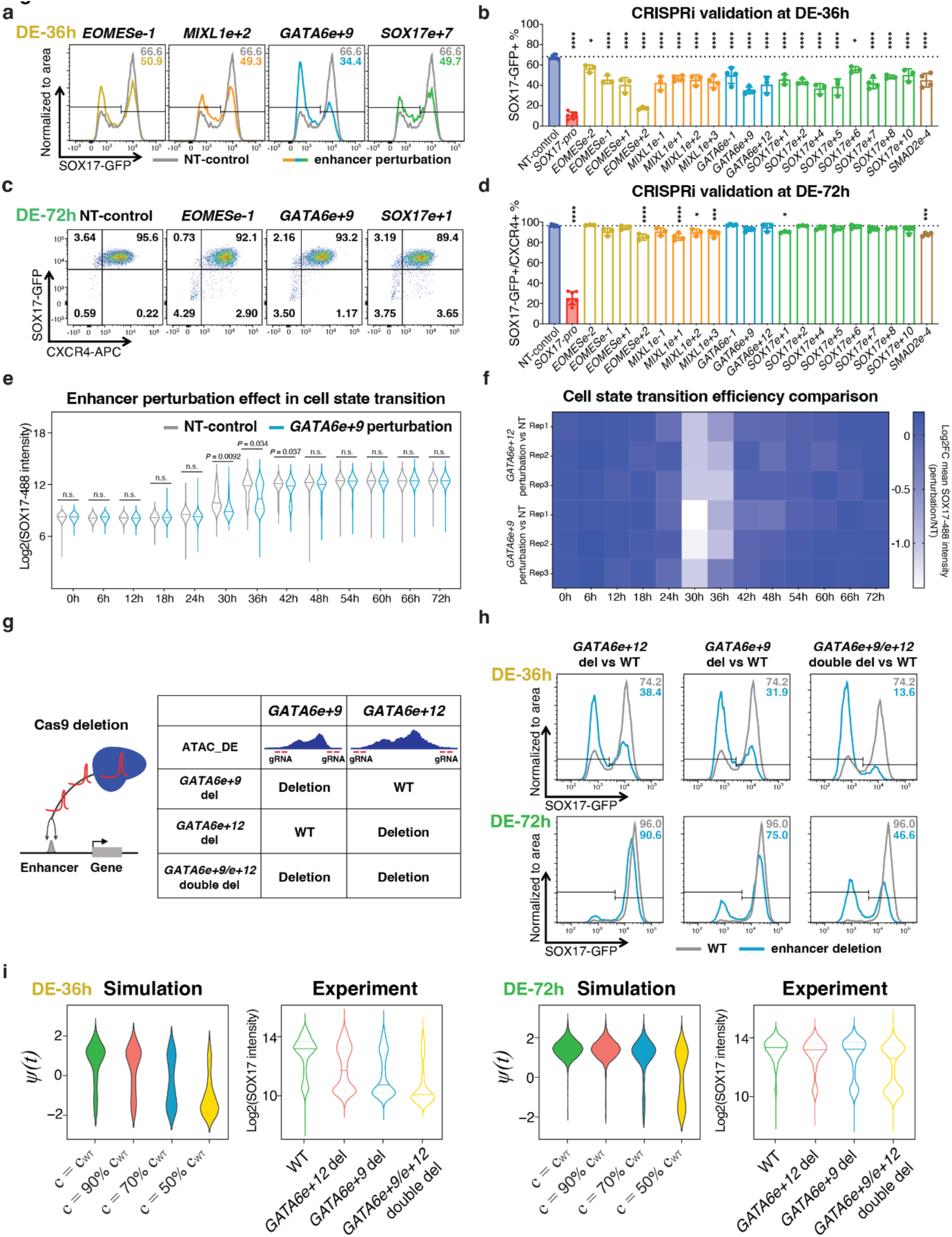
Validation of identified core enhancers using CRISPRi perturbation and CRISPR-Cas9 mediated deletion. **a-d**, Representative flow plots showing individual core enhancer perturbations decrease the hESC-DE transition efficiency measured by SOX17-GFP^+^ at DE-36h (**a**) and SOX17-GFP^+^/CXCR4-APC^+^ at DE-72h (**c**). The bar graphs show the percentage of SOX17-GFP^+^ cells at DE-36h (**b**) and SOX17-GFP^+^/CXCR4-APC^+^ at DE-72h (**d**). n≥3 independent experiments. Error bars indicate mean ± SD. Statistical analysis was performed by two-tailed unpaired multiple comparison test with Dunnett correction. Significance is indicated as: *P < 0.05, **P < 0.01, ***P < 0.001 and ****P < 0.0001. **e**, Violin plots showing the effect of *GATA6e+9* perturbation on hESC-DE transition efficiency (SOX17 intensity) measured every 6h through flow cytometry. n=3 independent experiments. Solid lines indicate median. Dashed lines indicate quartiles. Statistical analysis was performed by two-tailed paired student t-test with mean of each replicate. n.s.: not significant. **f**, Heatmap showing the comparison between the mean SOX17 expression intensity of cells with non-targeting control (NT) vs *GATA6e+9* or *GATA6e+12* perturbation measured every 6h through flow cytometry during hESC-DE transition. A significant impact was observed only in a narrow time window (around DE-36h) during the transition. **g**, Schematics of *GATA6e+9* del, *GATA6e+12* del and *GATA6e+9/e+12* double del hESC lines generation using two pairs of gRNAs with Cas9. **h**, Flow plots showing *GATA6* core enhancer del reduced hESC-DE transition efficiency at DE-36h and DE-72h. **i**, The comparison between the simulation and experiment results of the impact of different levels of enhancer perturbation on cell state transition. Solid lines indicate median. Dashed lines indicate quartiles.

We reasoned that the GRN is robust in part because multiple enhancers could interact additively or synergistically at a locus to regulate target gene expression. We applied CRISPR-Cas9 to generate single and double deletions of *GATA6e+9* and *GATA6e+12* (Fig.3g; Extended Data Fig.5c). The deletion of both *GATA6* enhancers showed a stronger impact on transition efficiencies compared to the expected additive effects of individual enhancer deletions at both DE-36h and DE-72h (Fig.3h; Extended Data Fig.5d, e). These results support synergistic interactions of these two core *GATA6* enhancers. Compared to single enhancer deletions, the double deletion produced a clearer separation of differentiated and undifferentiated cells at DE-72h. These experimental data closely match results from the dynamic GRN modeling (Fig.3i). We also compared the enhancer deletion results with single or double perturbations using CRISPRi and found that the latter had mild or no impact at DE-72h (Extended Data Fig.5d). Thus, although CRISPRi is more amenable for large-scale enhancer screens than CRISPR-Cas9 mediated deletion, it may miss *bona fide* enhancers especially when examining the perturbation effects in a steady state. The weaker effect of dCas9-KRAB could be due to competition with endogenous TFs and other technical differences between CRISPRi and deletion^38^. Overall, our findings demonstrate that the GRN is robust post-transition but can still be disrupted with strong perturbations as achieved here through the double deletion of *GATA6* enhancers.

## CTCF loop-constrained Interaction Activity (CIA) model improves enhancer prediction

Our CRISPRi screen and validation studies identified multiple enhancers around the core DE regulators (GATA6, EOMES, SOX17, and MIXL1; Extended Data table 4). This high-quality dataset allowed us to explore genomic features that can distinguish the enhancers which were positive in the screen (hits) from those which were not (non-hits). As expected, all enhancer hits for the core DE TFs had elevated levels of H3K27ac and increased accessibility during hESC-DE transition, accompanied by the binding of DE core TFs EOMES, GATA6 and SOX17 (Extended Data Fig.6). However, we also noticed that the hits were often located on one side of the TSS. We performed Hi-C and CTCF ChIP-seq experiments in ESC and DE, and observed strong concordance among bounded domains of increased Hi-C contact frequency (often referred to as topologically associated domains, or TADs), CTCF loops measured by ChIA-PET in H1 hESCs ^39^, CTCF loops predicted by our loop competition and extrusion model (LE model) ^40^ and CTCF binding (Fig.4a). While Hi-C does detect increased interactions between the promoter and distal enhancer hits (i.e., *SOX17e+10*) in DE, Hi-C contact frequency between the promoter and distal enhancers is only weakly correlated with the effect of enhancer perturbation (Fig.4c). In addition, enhancer hits are often distributed broadly around the target gene (Fig.2h; Extended Data Fig.4f), while the Hi-C contact signals are primarily enriched near the promoter (Fig.4a). In contrast, all the *SOX17* enhancer hits fall into a CTCF loop enclosing the promoter (Fig.4a), both as measured by CTCF ChIA-PET ^39^ and as predicted by the LE model ^40^. To quantify, we calculated the Hi-C contact frequency (with the promoter) for each enhancer along the locus. We also computed the probability (denoted as *P*(in loop)) that each distal enhancer is enclosed within the same CTCF loop as the promoter, using the ratio of the sum of counts for all loops enclosing both enhancer and promoter to the sum of all counts for all loops enclosing the promoter (Methods). Comparing the genomic intervals with large *P*(in loop) against those intervals with large Hi-C contact frequency, the former is broader and encompasses more hits (Fig.4a). Similar observations hold for the other loci (Extended Data Fig.7a). This suggests that we are less likely to miss impactful enhancers based on predictions by their enclosure within a CTCF loop than by enhancer-promoter Hi-C contact frequency.

To quantitatively compare the predictive power of these chromatin conformation features individually, we plot precision-recall curves for predicting hits (log2FC > 0.15) from non-hits for all DE gene enhancers (Fig.4b; Extended Data Fig.7b; Extended Data table 4). *P*(in loop) is more predictive (AUPRC=.818) than Hi-C contact frequency in DE (.692), Hi-C contact frequency in ESC (.598) or 1/|distance| from the promoter (.604). Many non-hit enhancers with a high *P*(in loop) have low chromatin accessibility, TF binding, and H3K27ac (Fig.4a; Extended Data Fig.6). Therefore, we tested all combinations of these features and the enhancer-promoter interaction information based on CTCF looping or Hi-C, using logistic regression, and assessed both AUPRC and correlation with log2FC (Fig.4c, d). Among the potential enhancers we tested, enclosure within a promoter-containing CTCF loop is more predictive than any other single feature. Combining interaction information with ATAC, H3K27ac and TF binding improves AUPRC, and in all cases CTCF loop-based models are more predictive than Hi-C contact frequency-based models. The combination of CTCF loop, ATAC, and H3K27ac achieves a very accurate prediction with AUPRC=.898. There is a further slight improvement by adding core TF binding data from EOMES, GATA6, SOX17 ChIP-seq (AUPRC=.925). Adding H3K4me1 to ATAC and H3K27ac can slightly improve the predictive power, but the prediction with H3K4me1 is not as good as H3K27ac when either is combined with ATAC alone or ATAC and core TF binding (Extended Data Fig.7c). The ABC model ^4^ combines ATAC and H3K27ac into 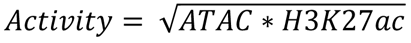, and uses either Hi-C contact frequency or 1/|distance| as contact measurement. We found that CTCF loop-constrained models outperformed both (Fig.4c, d). Nevertheless, we found that combining the 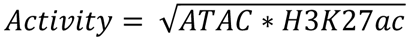 with CTCF loop information is simpler and performs comparably to logistic regression using ATAC and H3K27ac independently (Extended Data Fig.7d). The combination of *P*(in loop) > 0.5 and can classify most hits more cleanly than in combination with Hi-C (Extended Data Fig.7e). Our best classification result is with both *P*(in loop) above a threshold near 0.5 and 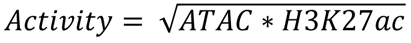greater than a threshold value near 1; as shown in Fig.4e, almost all hits (green) are correctly classified, and non-hits are correctly classified either by being outside a CTCF loop (grey) or below the *Activity* threshold curve (red). Since both *P*(in loop) and *Activity* are required, we will refer to the combined rule *CIAscore* = *P*(in loop) ∗ as the CTCF loop-constrained Interaction Activity (CIA) model.

We further evaluated the generalizability of this model with additional CRISPRi datasets from K562 cells in Reilley *et al* ^9^ and Nasser *et al* ^41^. The latter study summarized screening data from multiple sources ^4, 6, 11–15^. After mapping gRNAs to distal accessible regions (Dnase hypersensitive sites, or DHS peaks) in K562 cells, we identified 36 hits and 414 non-hits around 12 genes from Reilley *et al* (Extended Data Fig.8a; Extended Data table 5) and 69 hits and 1862 non-hits from Nasser *et al* (Extended Data Fig.9a; Extended Data table 6). Using these data, we also found that CTCF loop information was more predictive than Hi-C contact frequency (Extended Data Fig.8b; Extended Data Fig.9b), and CTCF loop + *Activity* was more predictive than the ABC model or Hi-C + *Activity* (Extended Data Fig.8c, d; Extended Data Fig.9c, d). An *Activity* threshold again clearly distinguished hits within loops (Extended Data Fig.8e; Extended Data Fig.9e). To better compare between the CIA and ABC models, we integrated all three datasets (ours, Reilley et al., and Nasser et al.). We scaled log2FC to “effect size” (Methods) and show the effect size vs. distance from TSS for all enhancer-promoter pairs tested (Fig.4f). For an equal number of positive predictions by CIA and ABC in each dataset (24+36+69, fixed recall), many strong effect enhancers are predicted by both models (yellow). However, the predictions made by CIA alone (red) have larger effects than predictions made by ABC alone (blue). Adding TF binding to the CIA model more clearly distinguishes hits from non-hits and performs slightly better (Fig.4d). In the absence of TF binding information, gkm-SVM scores can reproduce this small improvement in predictive accuracy (Extended Data Fig.7d; Extended Data Fig.8c; Extended Data Fig.9c). In summary, using three large functional enhancer screening datasets, we show that enhancer impact is more reliably predicted by enhancer location within a CTCF loop than by direct enhancer-promoter Hi-C contact frequency, and we constructed a simple predictive model of enhancer impact by combining CTCF loop-constrained interaction and enhancer activity.

## Discussion

We performed a CRISPRi screen for enhancers controlling the stimulated hESC-DE transition and discovered many novel enhancers flanking each core DE TF gene. ChIP-seq results show that each of these enhancers are bound by multiple core TFs, supporting that cooperative autoregulation through multiple TFs at enhancers is a prevalent feature of GRNs controlling cell state. All top enhancers were successfully validated individually. The results revealed that their perturbation delays the transition to DE, and the effect of enhancer perturbation is almost negligible after the transition to DE has been completed. A similar delayed phenotype in response to enhancer perturbation has been observed at the *Hox* cluster in flies ^42^ and mice ^43, 44^. Simulations of the hESC-DE transition using our dynamic GRN model agree with the observation of a delayed transcriptional response to enhancer perturbation, and show that nonlinear saturation of enhancer activity post-transition is responsible for this robustness once the GRN has fully activated the core TFs. In our simulations, the hysteresis of the GRN occurs via autoregulatory enhancer activity coupled to the translation of the TFs, and is at a longer time scale than simulations of nonlinearities which may also be present at the time scale of transcriptional activation ^45^. Together, these observations provide a plausible explanation for why variants and putative enhancer regions from genome-wide association studies (GWAS) are often difficult to validate when tested individually in the steady state. These enhancers may contribute to cell state transitions and cell abundance while not strongly affecting transcript levels in the established state or when tested in isolation. This suggests a direct mechanism by which enhancers may contribute to human disease even in the absence of strong effects in post-transition cells, and recent reports have emphasized that cell type abundance quantitative trait loci (QTLs) can have profound impact on human phenotypes ^46–48^. Our results also predict that screens designed to target enhancers during a cell state transition will have greater sensitivity of detection. In addition, one may explore ways to weaken the GRN (e.g., through manipulating a core TF or enhancer) to increase its vulnerability to further enhancer perturbation, as suggested by the *GATA6* enhancer double deletion studies. Such strategies could mitigate the challenge of enhancer redundancies and support a more comprehensive discovery of enhancers. Beyond developmental cell fate changes, we envision that stimulating physiological or pathological cell state transitions can accelerate the discovery of disease-relevant enhancers or variants.

The sensitivity of our screen also revealed that most functional enhancers fell within CTCF loops, leading us to propose an interaction model whereby CTCF loops constrain enhancer interactions and activity (CIA model). Our large number of validated enhancers allowed comparisons between alternative hypotheses and showed that the CTCF constraint is significantly more predictive than Hi-C-based measurements of contact frequency between the enhancer and target promoter. Many distal enhancer hits within CTCF loops have large transcriptional impact but relatively low enhancer-promoter contact frequency as measured by Hi-C. Adjusting for the distance-dependency in Hi-C data through simple power law distance corrections did not significantly improve AUPRC (Methods), but it may be possible to improve the predictions through developing newer methods based on Hi-C data or combining Hi-C with CTCF and other datasets. Mechanistically, while the prevailing model supports that transcriptional activity depends on enhancer-promoter contact, our observation at multiple enhancers implies a highly nonlinear relationship between contact frequency and transcription. This notion is supported by recent studies suggesting that the contact probability has a nonlinear impact on transcription ^45, 49^, and a contact-independent model has also been proposed ^50^. Through simulations, multiple plausible mechanisms have been shown to generate this nonlinear relationship, including accumulation of promoter bound factors or post transcriptional modifications at the promoter ^45^. Instead of relying on direct enhancer-promoter contact frequency measurements from Hi-C data, our CIA model utilizes CTCF loop information to constrain enhancer prediction. We combined the CTCF constraint with *Activity* and showed that this CIA model outperforms the ABC model on results from our screen and screens conducted in K562 cells from other groups ^9, 41^ This simple CIA model should be quite useful for prioritizing and understanding single nucleotide polymorphisms (SNPs) implicated in expression QTLs (eQTLs) or GWAS-associated loci.

Our CIA model is a quantitative improvement over the “insulated neighborhood” hypothesis ^51^ and is consistent with many studies showing that enhancer activity is largely restricted within CTCF loops and TADs ^49, 52, 53^, and that TAD disruption can lead to oncogenic misexpression and developmental diseases ^54, 55^. The CTCF loops are visually consistent with Hi-C TADs. However, TAD calling provides a binary result of whether a genomic region is within a TAD, which can be unstable and sensitive to parameters and methods. In comparison, *P*(in loop) calculates the probability that a genomic region is enclosed within a CTCF loop, thus providing a more reliable measurement. However, there are also seemingly conflicting findings that auxin degradation of CTCF can only have a modest effect on transcription on a short time scale ^56^. We speculate that during cell state transitions, CTCF-mediated enhancer-promoter proximity is necessary for establishing *de novo* enhancer-protein-promoter complexes (specific to the succeeding cell state). While in post-transition steady states, the established complex can reinforce the proximity together with the CTCF loop. Without the CTCF loop, the proximity can still be maintained, though it will likely become sub-stable, and the transcription efficiency may gradually decrease. Our dynamic GRN model shows a similar result that once the active transcriptional state is established, the transcriptional response to a perturbation of enhancer activity occurs at a much longer time scale (Fig.1g). The hysteresis in our model thus provides a plausible explanation for the long-standing paradox that CTCF is required to restrict enhancer activity, yet removing CTCF does not immediately affect transcription.

## Methods

### Cell lines and culture conditions

iCas9 *SOX17^eGFP/+^* HUES8 and idCas9-KRAB *SOX17^eGFP/+^* HUES8 hESCs were cultured on vitronectin-coated (Gibco; A14700) plates and maintained in E8 medium (Gibco; A1517001). Cells were dissociated by 3-5 mins treatment with 0.5 mM EDTA in 1X DPBS without calcium and magnesium at room temperature. 10 μM ROCK inhibitor Y-27632 (Selleck Chemicals; S1049) was added into E8 medium for the first 24h after passage. Medium was changed every day. hESCs were passaged every 3-4 days depending on the cell growth speed and confluency. All cell lines were routinely tested by Memorial Sloan Kettering Cancer Center (MSKCC) Antibody & Bioresource Core Facility to confirm there was no mycoplasma contamination and by MSKCC Molecular Cytogenetics Core to confirm there were no karyotyping abnormalities. Experiments with hESCs were conducted per NIH guidelines and approved by the Tri-SCI Embryonic Stem Cell Research Oversight (ESCRO) Committee.

### hESC definitive endoderm differentiation

We found that DE differentiation is more robust when using hESCs during the logarithmic growth phase. To optimize the differentiation, cells were maintained in the logarithmic phase before seeding for DE differentiation with a recovery passage as follows. Please note that the cell numbers described below were optimized based on the growth of the HUES8 hESCs in our laboratory, and they may need to be adjusted for other hESC lines. hESCs were dissociated with 1X TrypLE Select (Gibco; 12563029) for 3 mins at room temperature. Then TrypLE was removed, and cells were washed and resuspended into E8 medium with 10 μM ROCK inhibitor Y-27632. Cells were counted by using Vi-CELL XR Cell Viability Analyzer (Beckman Coulter), and 2-3 million cells were seeded per 10-cm plate with E8 medium containing 10 μM ROCK inhibitor Y-27632. The E8 medium was refreshed daily, and the cell number should reach 10-12 million per 10-cm plate 2 days after seeding. hESCs after the recovery passage were collected and counted again as described above and 1 million cells per well were seeded in a 6-well plate in E8 medium with 10 μM ROCK inhibitor Y-27632; and 18h later, hESCs were changed into fresh E8 medium. At 24h after seeding, hESCs should reach 2-2.4 million per well, and be ready for differentiation. For DE differentiation, hESCs were first washed with 1X DPBS once and then cultured in 2.5 ml S1/2 differentiation medium daily as previously described ^25, 57^: cells were treated with 50 ng/ml Activin A (Bon-Opus Biosciences; C687-1mg) for three days, and 5 µM CHIR99021 (TOCRIS; 4423) for the first day.

### scRNA-seq

scRNA-seq was performed as previously described ^58^. Cells were harvested every 12h during DE differentiation for scRNA-seq experiments with targeted collection ranging from 3000 to 6000 cells. Single-cell 3′ RNA-seq libraries were generated with 10x Genomics Chromium Single Cell 3′ Reagent Kit v.3 following the manufacturer’s guidelines. The libraries were sequenced on NovaSeq 6000 platform following the manufacturer’s guidelines.

### scRNA-seq PCA analysis

Chromium v3 analysis software Cell Ranger (Version 3.1.0) was run using “cellranger count – expect-cells 1000”. Genes with fewer than 10 reads summed over all cells were removed, yielding 21099 transcripts detected across 35988 cells, approximately 5000 cells per time point. The cells ranked in the bottom 10% of total transcript number per cell were removed from further analysis. For PCA we further restricted analysis to transcripts whose standard deviation across the 7 time points was greater than 0.2.

### scRNA-seq gene expression correlation analysis

Per best practices of Seurat package ^59^, the quality control of cells included the number of features found in a cell and the percentage of tags mapped to the mitochondrial chromosome. Each sample was filtered according to the following criteria:

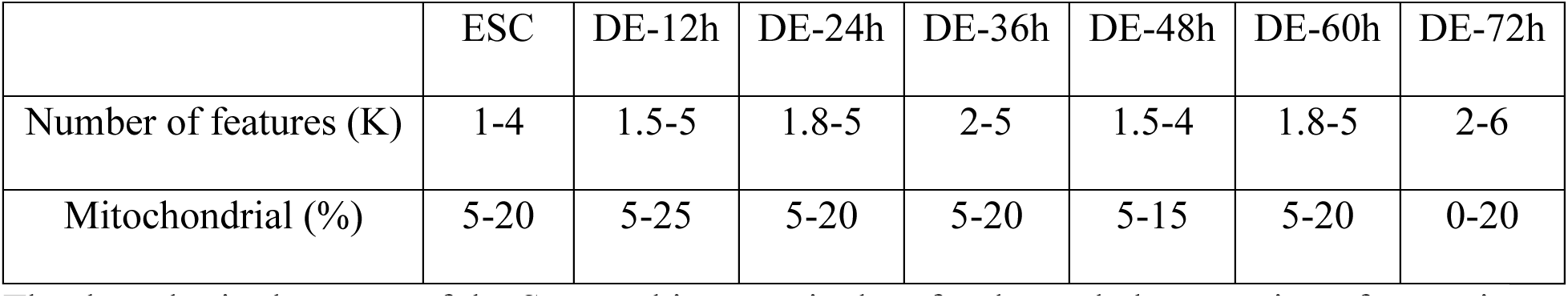

|  | ESC | DE-12h | DE-24h | DE-36h | DE-48h | DE-60h | DE-72h |
| --- | --- | --- | --- | --- | --- | --- | --- |
| Number of features (K) | 1-4 | 1.5-5 | 1.8-5 | 2-5 | 1.5-4 | 1.8-5 | 2-6 |
| Mitochondrial (%) | 5-20 | 5-25 | 5-20 | 5-20 | 5-15 | 5-20 | 0-20 |

The data slot in the assay of the Seurat object, equivalent for the scaled expression of genes in cells, was used for gene-wise visualization by UMAP. For selecting the relevant expressed genes, each gene was applied a filter where the percentage of nonzero expression in cells was at least 55 percent at any time point. We focused on relevant expressed genes by removing mitochondrial, ribosomal, miRNAs, lincRNAs, antisense transcripts, and genes that were not yet named (e.g., those containing orf and starting with AC or AL). The matrix from the 2606 relevant expressed genes, after transposition for gene-centric embedding, was embedded in UMAP using the R package umap, using the Pearson correlation as the distance metric of umap.config. Each point was labeled with the color scale defined from the expression fold change from ES to DE-72 hr.

### Gene Regulatory Network Model

The network state in our GRN model is described by a vector of core TF concentrations *ψ* = (*ψ*_!,_ *ψ*_2_, …, *ψ*_$_) = (*ψ*_34!..$_). For example, a network with three genes and *ψ* = (*A*, *B*, *C*) is shown in Fig.1a. Each component TF gene *i* is described by an equation similar to Eq (1):

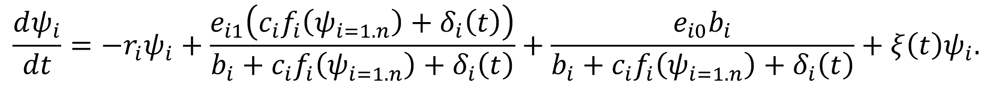

The activation probabilities for gene *i* are *p*_on_∼*c*_3_ *f*(*ψ*_*i=1.n*_) + *δ*_3_ (*t*) and *p*_oof_∼*b*_3_, so together the probability that gene *i* is activated and transcribed at rate 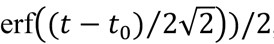and the probability that gene *i* is transcribed at basal rate 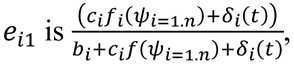, with parameters *b*_3_, *c*_3_, and *δ*_3_ (*t*) specific to each core TF in the network. Each term *f*_3_ (*ψ*_*i=1.n*_) will be a function of the activity of the core TFs binding gene *i*’s enhancers. As a consequence of strong nonlinear cooperativity, we now make the simplifying assumption that the core TFs turn on synchronously (*α*_!_*ψ*_!_ ≈ *α*_2_*ψ*_2_ ≈ ⋯ ≈ *α*_$_*ψ*_$_) where *α*_34!..$_ are constants. This allows us to approximately represent the entire network activity with the scalar *ψ*(*t*) and replace the gene specific enhancer activities with an average enhancer activity for each core TF gene, *f*_3_(*ψ*_*i=1.n*_) ≈ *cψ*^$^, where *n* now represents an average degree of cooperativity, and could arise from either individual TF cooperativity at an enhancer or from multiple enhancers interacting non-additively. Because the probability that the gene is activated quickly saturates at large *ψ*, the precise form of this nonlinearity does not strongly affect the results, as long as *n* ≥ 3. We similarly replace gene specific rates *b*_3_, *c*_3_, and *δ*_3_ (*t*) with weighted averages, leading to Eq. (1). Eq.1 was solved by the Euler-Maruyama method using parameters (*b*, *c*, *e*_"_, *e*_!_, *r*, *B*) = (.5,1, .1,3,1,3) unless stated otherwise, and with normally distributed noise *ξ*(*t*) with amplitude *ξ*_"_ = 0.4. For *e*_!_ ≫ *e*_"_ the OFF fixed point is at *ψ* ≈ *e*_"_/*r* and the ON fixed point is at *ψ* ≈ *e*_!_/*r* and both are stable for *cbr*/*e*_!_ and *B* ≥ 3. The stimulus *δ*(*t*) was modeled as *δ*(*t*) = *δ*_0_(1 + erfS(*t* − *t*_"_)⁄2√2T)/2, with *δ*_0_ = 0.035, which turns on smoothly at *t* = *t*_"_, to model the sustained effect of Activin A in the experiments, which is known to act through WNT/Nodal signaling. The stimulus produces a weak increase in enhancer activity (*p*_’(_), that is independent of *ψ*, and is therefore independent of core TF network activity. Although the bifurcation and stability results are valid for any *e*_!_ ≫ *e*_"_ and *cbr*/*e*_!_ and *B* ≥ 3, the parameters *δ*_0_, *ξ*_0_, *e*_0_, and *e*_0_ were chosen to approximately match the distributions of SOX17 expression levels and population variation in the FACS data. A model with a similar mathematical form was previously derived to model cellular differentiation induced by MAPK signaling and was also shown to admit bistable solutions ^60, 61^.

### Stochastic Gillespie simulations

The stochastic Gillespie algorithm was used to simulate a strongly nonlinear cooperative autoregulatory gene circuit model ^33, 34^, analogous to the rate equation Eq.1 but also valid for arbitrarily small numbers of molecules. In this model, up to three molecules of the TF *A* sequentially bind to an enhancer (*E*), and the differentially bound TF-enhancer complexes are denoted as *EA*, *EAA*, and *EAAA*. The eight possible reactions, their probabilities, *a*_3_, and their impact on molecular species numbers are given as follows, where *X*_6_ indicates the number of molecules of species *M*:

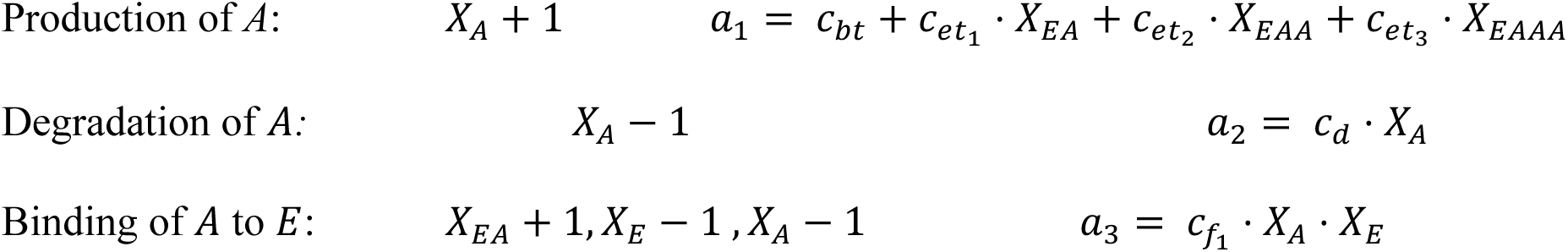
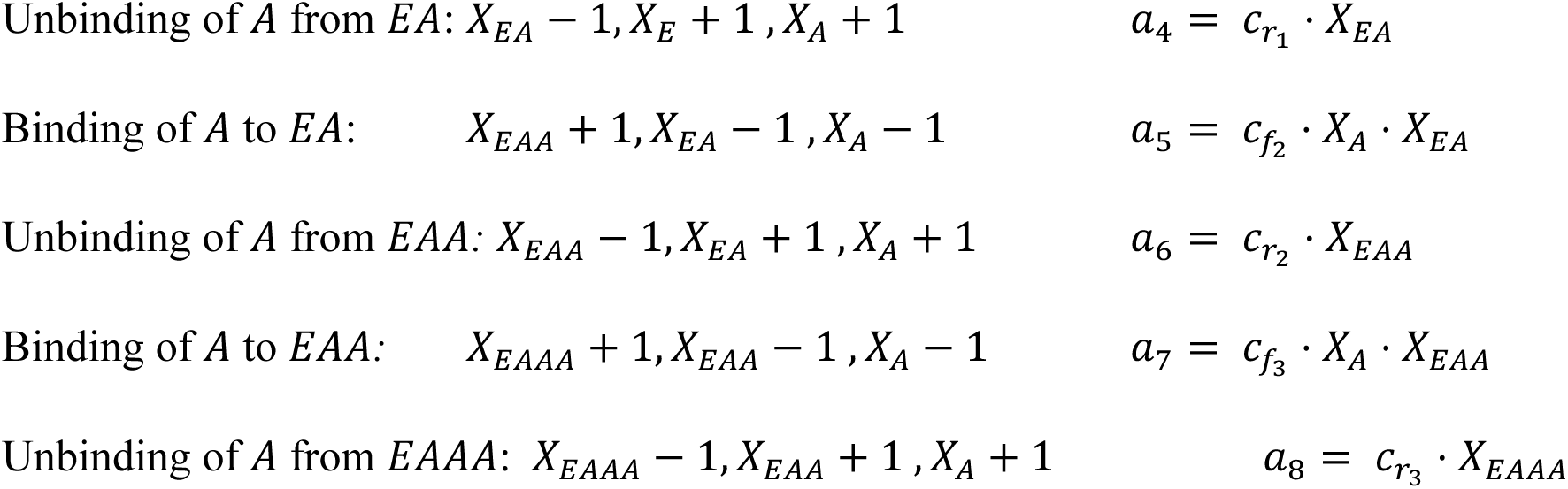

Unless noted otherwise, the rates used are S*c_bt_, c_et1_, c_et1_, c_f1_, c_r1_, c_f2_, c_r2_, c_f3_, c_r3_, c_d_, c_d_, T = (. 04, .04, .04, 2.4, .1, 15, .1, 15, .1, 15, .003). The strong cooperativity of enhancer activity is reflected in the fact that we chose *c*_)(9_ ≫ *c*_1(_, *c*_)(!_, *c*_)(2_. Fifty independent runs, each representing a single cell, were performed for each datapoint shown. The initial enhancer state at *t* = 0 is given by (*X_E_, X_EA_, X_EAA_, X*_EAAA_*) = (1,0,0,0), and the initial number of TFs, *X_a_*, was sampled from a uniform distribution. The stimulus is modeled by a transitory increase in the initial rate of *A* binding to *E*, *c_f1_*.

### Generation of the idCas9-KRAB *SOX17^eGFP/+^* HUES8 hESC line

idCas9-KRAB *SOX17^eGFP/+^* HUES8 hESC line was generated using a cassette switch strategy based on previously established iCas9 *SOX17^eGFP/+^* HUES8 hESC line ^25, 62^. iCas9 *SOX17^eGFP/+^* HUES8 hESCs were treated with 10 μM ROCK inhibitor Y-27632 and 2 μg/ml doxycycline one day before transfection. gRNAs that specifically targeted on *AAVS1-iCas9* allele were co-transfected with AAVS1-idCas9-KRAB donor vector into the cells by using Lipofectamine Stem Transfection Reagent (Thermo Fisher Scientific; STEM00001) following manufacturer’s guidelines. Transfected cells were treated by Hygromycin selection for 7 days and single cell colonies were picked for genotyping. Inducible dCas9-KRAB were validated by RT-qPCR and flow cytometry analysis. gRNAs and genotyping primer sequences are listed in Extended Data table 7-8.

### ChIP-MS and analysis

ChIP-MS and analysis were performed as previously described ^63^ with minor modifications. Antibodies used for immunoprecipitation are listed in Extended Data table 9. Briefly, ChIP-MS was performed using the same ChIP protocol as in ChIP-seq. Cells were crosslinked with 1% formaldehyde and then lysed and sonicated. Clear supernatant was collected for chromatin immunoprecipitation. Total protein was eluted after immunoprecipitation by incubation with 5% SDS and 5mM DTT at 98°C for 5 min. Then protein samples were alkylated, trypsinized, and desalted for LC-MS/MS acquisition. Each experimental condition was performed with two biological replicates. Protein levels were log2 transformed, normalized by the average value of each sample and missing values were imputed using a normal distribution 2 standard deviations lower than the mean before statistics analysis. Protein levels from EOMES, GATA6 and SOX17 ChIP-MS were further compared to IgG controls and the common interacting TFs with log2FC > 2 and -log10(p-value) > 2 were used to plot Fig.2d.

### RNA isolation, reverse transcription, and real-time quantitative PCR (RT-qPCR)

Total RNA was extracted using Quick-RNA MiniPrep kits (ZYMO research; R1055) following the manufacturer’s guidelines. cDNA was produced by using High Capacity cDNA Reverse Transcription Kit (Applied Biosystems; 4368817) with 2 μg of total RNA per reaction. RT-qPCR reaction was performed with SYBR Green Master mix (Applied Biosystems; A25742) in the 7500 or QuantStudio 6 flex Real Time PCR system (Applied Biosystems). GAPDH was used as an internal control. Primers used in RT-qPCR are listed in Extended Data table 8.

### RNA-seq

After RiboGreen quantification and quality control by Agilent BioAnalyzer, 500ng of total RNA with RIN values of 6.5-10 underwent polyA selection and TruSeq library preparation according to instructions provided by Illumina (TruSeq Stranded mRNA LT Kit; RS-122-2102), with 8 cycles of PCR. Samples were barcoded and run on a HiSeq 4000 or NovaSeq 6000 platform in a PE50 run, using the HiSeq 3000/4000 SBS Kit or NovaSeq 6000 SP or S2 Reagent Kit (100 Cycles) (Illumina).

### RNA-seq analysis

We followed the ENCODE RNA-seq processing pipeline, aligning reads to hg38 with STAR_2.5.1b and parameters “--outFilterMultimapNmax 20 --alignSJoverhangMin 8 -- alignSJDBoverhangMin 1 --outFilterMismatchNmax 999 --outFilterMismatchNoverReadLmax 0.04 --alignIntronMin 20 --alignIntronMax 1000000 --alignMatesGapMax 1000000”. Transcripts were quantified with RSEM v1.2.23.

### ATAC-seq

Profiling of chromatin was performed by ATAC-Seq as previously described ^64^. Briefly, 50K cryopreserved cells were washed in cold PBS and lysed. The transposition reaction containing TDE1 Tagment DNA Enzyme (Illumina; 20034198) was incubated at 37°C for 30 minutes. The DNA was purified with the MinElute PCR Purification Kit (QIAGEN; 28004) and amplified for 5 cycles using NEBNext High-Fidelity 2X PCR Master Mix (New England Biolabs; M0541L). After evaluation by real-time PCR, 3-14 additional PCR cycles were done. The final product was cleaned by AMPure XP beads (Beckman Coulter; A63882) at a 1X ratio, and size selection was performed at a 0.5X ratio. Libraries were sequenced on a HiSeq 4000 or NovaSeq 6000 platform in a PE50 run, using the HiSeq 3000/4000 SBS Kit or NovaSeq 6000 S1 Reagent Kit (100 Cycles) (Illumina).

### ATAC-seq analysis

Paired-end reads were mapped to hg38 with bowtie2 using default parameters, duplicate reads were removed with picard, and peaks were called using macs2 using default parameters. gkm-SVM was run on top 10000 300bp distal peaks (>2 kb from TSS) and negative sequence following ^24, 31, 32^. Motifs were extracted using gkm-PWM.

### gRNA design of core enhancer perturbation screen

2 Mb up/down stream of the 10 core TFs were selected for further chromatin accessibility filtering. Only the regions showing accessibility in either ESC stage or DE-48h stage were kept. Regions that overlapped with promoters or exons were removed from the list, resulting in 394 putative enhancers being selected in total. gRNA design was performed by using CHOPCHOP ^55^ to achieve full-tiled coverage of the selected regions. We further added 3 gRNAs targeting *SOX17* promoter, and 1,100 gRNAs targeting safe harbor loci ^37^ as positive and negative controls, respectively. gRNA sequences of the library are listed in Extended Data table 2.

### Oligo synthesis and library cloning

gRNA oligos were synthesized on-Chip (Agilent). Synthesized oligos were amplified and restriction cloned into lentiGuide-puro (Addgene; 52963) by the MSKCC Gene Editing & Screening Core Facility. Cloned plasmid library was PCR amplified to incorporate adapters for NGS. Samples were purified and sequenced using Illumina HiSeq 2500 platform. FASTQ files were clipped by position and reads were mapped back to the reference library file to show relative abundance of reads per gRNA. Reads within each sample were normalized to total number of mapped reads and library size. The Overall Representation of Library was charted over a one-log fold change to evaluate if any gRNA was over-or under-represented in the final library. Primers used for PCR are listed in Extended Data table 8.

### Lentiviral library generation

The core enhancer perturbation lentiviral library generation was performed as previously described ^25^ with minor modifications. Briefly, a total of 13.6 μg core enhancer perturbation library plasmids with 5.44 μg lentiviral packaging vector psPAX2 and 1.36 μg vesicular stomatitis virus G (VSV-G) envelope expressing plasmid pMD2.G (Addgene plasmid 12260 and 12259) were transfected with the JetPRIME (VMR; 89137972) reagent into 293T cells to produce the lentivirus. Fresh medium was changed 24h after transfection and viral supernatant was collected, filtered, and stored at −80°C 72h after transfection.

### Core enhancer perturbation screen

We aimed for a ∼1,000-fold coverage per gRNA to maximize sensitivity. 35 million idCas9-KRAB *SOX17^eGFP/+^* HUES8 hESCs were collected and infected with the lentiviral library at a low MOI of ∼0.3 on Day 0 in 15-cm plates. A total of 6 μg/ml protamine sulfate per plate was added during the first 24h of infection to improve the infection efficiency. One day after infection (Day 1), cells were treated with 2 μg/ml doxycycline to induce dCas9-KRAB expression, which continues till the end of the screen at DE-36h. Infected cells were selected with 1 μg/ml puromycin from Day 2-Day 4 and harvested on Day 5 for recovery passage. 2 days after recovery passage, 60 million cells were collected and seeded into 15-cm plates for DE differentiation as described above. 36h after differentiation, cells were dissociated using 1X TrypLE Select and sorted using FACS Aria according to GFP expression. Cells whose GFP expression levels were in the top or bottom 20% were pelleted individually, with each pellet containing 15 million cells.

### gRNA enrichment sequencing and data analysis

The gRNA enrichment sequencing and data analysis were performed as previously described ^25^ with minor modifications, manipulated by MSKCC Gene Editing & Screening Core Facility. Briefly, genomic DNA from sorted cell pellets was extracted using the QIAGEN Blood & Cell Culture DNA Maxi Kit (QIAGEN; 13362) and quantified by Qubit (ThermoScientific; Q32850) following the manufacturer’s guidelines. A quantity of gDNA covering 1000X representation of gRNAs was PCR amplified to add Illumina adapters and multiplexing barcodes. Primer sequences to amplify lentiGuide-puro are shown in Extended Data table 8. Amplicons were quantified by Qubit and Bioanalyzer (Agilent) and sequenced on the Illumina HiSeq 2500 platform. Sequencing reads were aligned to the gRNA library sequences and counts were obtained for each gRNA. The read counts were normalized to total reads of each sample to offset differences in read depth. To calculate the Z-score of each gRNA, we subtracted the mean log2FC of all negative control gRNAs targeting safe harbors from the log2FC of each gRNA, and then divided the result by the standard deviation of log2FC from the negative control gRNAs (Extended Data table 2). Off-targets of each gRNA were further assessed by CRISPOR ^65^. gRNAs with 0 mismatch (MM)=1, 1MM<10, 2MM<30, 3MM<100 and total raw reads > 500 were kept for calculating average Z-score of each putative enhancer region. Putative enhancer region with less than 3 qualified gRNAs were filtered away (Extended Data table 3).

### Hit validation

The gRNA enrichment sequencing and data analysis were performed as previously described ^25^ with minor modifications. Briefly, selected gRNAs for each enhancer hit were cloned into lentiGuide-puro (for single perturbations) and lentiGuide-blast (when a second perturbation was used in combination). 1.36 μg lentiGuide-puro (or lentiGuide-blast), 0.1 μg pMD2.G and 0.4 μg psPAX2 plasmids were transfected with the JetPRIME (89137972; VMR) reagent into 293T cells to pack lentiviruses. Viral supernatant was made and collected as described above. idCas9-KRAB SOX17^eGFP/+^ HUES8 hESCs were then infected with viruses containing different gRNAs individually following the same process as described above for the screen. One day after infection (Day 1), cells were treated with 2 μg/ml doxycycline to induce dCas9-KRAB expression, which continues till the end of the experiment (DE-36h or DE-72h). Infected cells were selected with 1 μg/ml puromycin from Day 2-Day 4 and harvested on Day 5 for recovery passage and followed by DE differentiation described above. For dual selection, cells were selected with 1 μg/ml puromycin and 10 μg/ml blasticidin together for 5 days. Cells were collected at both 36h and 72h for flow cytometry analysis. gRNA sequences selected from the core enhancer validation are listed in Extended Data table 10.

### Flow cytometry

Flow cytometry analysis were performed as previously described ^25^. Antibodies used for flow cytometry are listed in Extended Data table 9. Briefly, for live GFP and surface marker data collection, cells were dissociated and stained with DAPI and corresponding antibodies. For TF data collection, cells were first stained with LIVE-DEAD Fixable Violet Dead Cell Stain (Invitrogen; L34955) and then fixed and stained with corresponding antibodies. Flow cytometry data were collected using BD LSRFortessa or BD LSRII. Flow cytometry analysis and figures were generated in FlowJo v10.

### Generation of clonal enhancer KO hESC lines

Enhancer KO hESC lines were generated by using two paired crRNAs surrounding targeted enhancers to increase knockout efficiency. crRNAs and tracrRNA were ordered from IDT. iCas9 *SOX17^eGFP/+^* HUES8 hESCs were treated with 2 μg/ml doxycycline and 10 μM ROCK inhibitor Y-27632 for 24h, dissociated with 1X TrypLE Select, and transfected with 0.15 μM of each crRNA and 0.6 μM of tracrRNA by using Lipofectamine RNAiMAX Transfection Reagent (ThermoScientific; 13778100) following the manufacturer’s guidelines. Transfected cells were further cultured in E8 with 10 μM ROCK inhibitor Y-27632 for 48h and ∼2000 cells were seeded into 100-mm plate to raise colonies. Then, the genomic DNA of individual colony was extracted by using DNeasy Blood & Tissue DNA Kit (QIAGEN; 69506) for genotyping. crRNA and genotyping primer sequences are listed in Extended Data table 7-8.

### ChIP-seq

ChIP-seq was performed as previously described ^25^ with minor modifications. Antibodies used for immunoprecipitation are listed in Extended Data table 9. Briefly, cells were crosslinked with 1% formaldehyde and then lysed and sonicated. Clear supernatant was collected for chromatin immunoprecipitation, decrosslinking and DNA purification. Then the sequencing library was generated by using the NEBNext® Ultra II DNA Library Prep Kit (New England Biolabs; E7103S) and NEBNext® Multiplex Oligos for Illumina® (New England Biolabs Index Primers Set 1; NEB, E7335S). Samples were pooled and submitted to MSKCC Integrated Genomics Operation core for quality control and sequencing on Illumina HiSeq 4000 platform.

### ChIP-seq/ChIA-PET analysis

Paired-end reads were mapped to hg38 with bowtie2 using default parameters, and peaks were called using macs2 using default parameters. Processed ChIA-PET bedpe files for K562, GM12878 ^66^, H1, and HUVEC ^39^ were downloaded from encodeproject.org.

### Hi-C

2 million cells were collected and fixed with 1% formaldehyde. The subsequent steps of Hi-C were then performed using the Arima-HiC kit (Arima; A510008) while libraries for sequencing were prepared with the KAPA Hyper Prep Kit (KAPA; KK8502) following the manufacturers’ guidelines. Samples were pooled and submitted to MSKCC Integrated Genomics Operation core for quality control and sequencing on Illumina HiSeq 4000 platform.

### Hi-C analysis

Prior to alignment, Hi-C Pro 2.11.4 was used to fragment, with GATC and GANTC as the restriction sites, with all alternative haplotypes removed. Reads were aligned with default settings for Bowtie 2.4.1 in Hi-C Pro, for both the global and local alignment steps. After both alignment steps, reads with a MAPQ of at least 30 were retained for further analysis and duplicates were removed. Sample level Hi-C maps were converted to .hic file format with JuicerTools 1.22.01. Condition specific Hi-C maps were generated by combining all sample level .allValidPairs files, and then converting them to the .hic file format with JuicerTools. Hi-C datafile ENCFF080DPJ.hic ^56^ for K562 was downloaded from encodeproject.org. Hi-C .hic datafiles for HUES64 and differentiated to (EC, MS, EN) were downloaded from NCBI GEO accession GSE130085 ^67^. Contact frequency .bedpe files were generated from Juicer Tools version 1.22.01 or 2.13.05 (K562).

### Predictive modeling of screen hits

Scores for ATAC, DHS, ChIP-seq, and H3K27ac signal features were mapped onto uniform 1000 bins centered on each target enhancer. For discriminative AUPRC analysis, positive hits were defined as log2FC > 0.15 for hESC-DE (24 positive hits out of 160 DE enhancers tested, Extended Data table 4). For K562 Reilly we downloaded all “Flow-Fish CRISPR Screen” “tsv” or “tsv guide quantification” files from https://www.encodeproject.org which yielded experiments at 20 loci^9^. For analysis of the Reilly data, we mapped gRNAs to non-promoter (>2 kb from TSS) K562 DHS peaks and normalized high and low expression gRNA counts. 12 genes had an enhancer hit with log2FC>0.8 (36 positive hits out of 450 K562 regions, Extended Data table 5) and we included these loci in model evaluation (Extended Data Fig 8a). For the Nasser data we downloaded Supplemental Table 6 from Nasser 2021^41^ and mapped all tested regions to non-promoter K562 DHS peaks within 1 Mb of a tested gene. Hits were defined by Regulated=TRUE in their Table 5 (69 positive hits out of 1931 K562 regions, Extended Data table 6), flanking 65 genes. For modeling, Hi-C contact frequency was calculated in 5kb bins from the .hic files normalized to one for the most promoter proximal bin using our data in ESC and DE, and in ENCFF080DPJ in K562. We also tried as a feature a distance corrected promoter Hi-C contact frequency, Hi-C * |dist|*^k^*, but found no improvement in AUPRC over *k*=0 for *k* between 0 and 2. *P*(in loop) for each enhancer-promoter pair was calculated from ChIA-PET reads for all loops in a 2 Mb window spanning the promoter. Total *P*(in loop) for each distal enhancer-promoter pair is given by the ratio of total ChIA-PET read counts of all loops spanning both the enhancer and the promoter divided by the total counts of all loops containing the promoter (but not necessarily also containing the enhancer). The minimum threshold for loop calls in the ChIA-PET data is either 3 or 4 reads, and we also used this threshold for loop counts. To reduce variability in the ChIA-PET data, we averaged counts for multiple datasets. For hESC-DE, H1, HUVEC, and GM12878 ChIA-PET was used, while for K562 Reilly and Nasser, K562 ChIA-PET was used, but different ChIA-PET datasets yield very similar *P*(in loop) and predictive performance. *P*(in loop) was normalized to one for the most promoter proximal bin. Logistic regression was used to combine features and predict performance. For these very low dimensional logistic regression models, test set performance is reduced by <2% compared to using the full dataset, which we used to reduce statistical variation when comparing all models. Spearman correlation was calculated between probability of being in the positive class and enhancer effect (log2FC). To compare all models in Fig 4f, log2FC in ESC-DE and K562 Reilly were scaled to “effect size” by dividing log2FC by 1.5 for ESC-DE and 8.0 for K562 Reilly so all datasets had a similar effect size threshold of 0.1 for hits. We took the top predictions from each model in each model to compare performance at constant recall (24 in ESC-DE, 36 in K562 Reilly, and 69 in K562 Nasser, 129 total).

**Figure 4.**
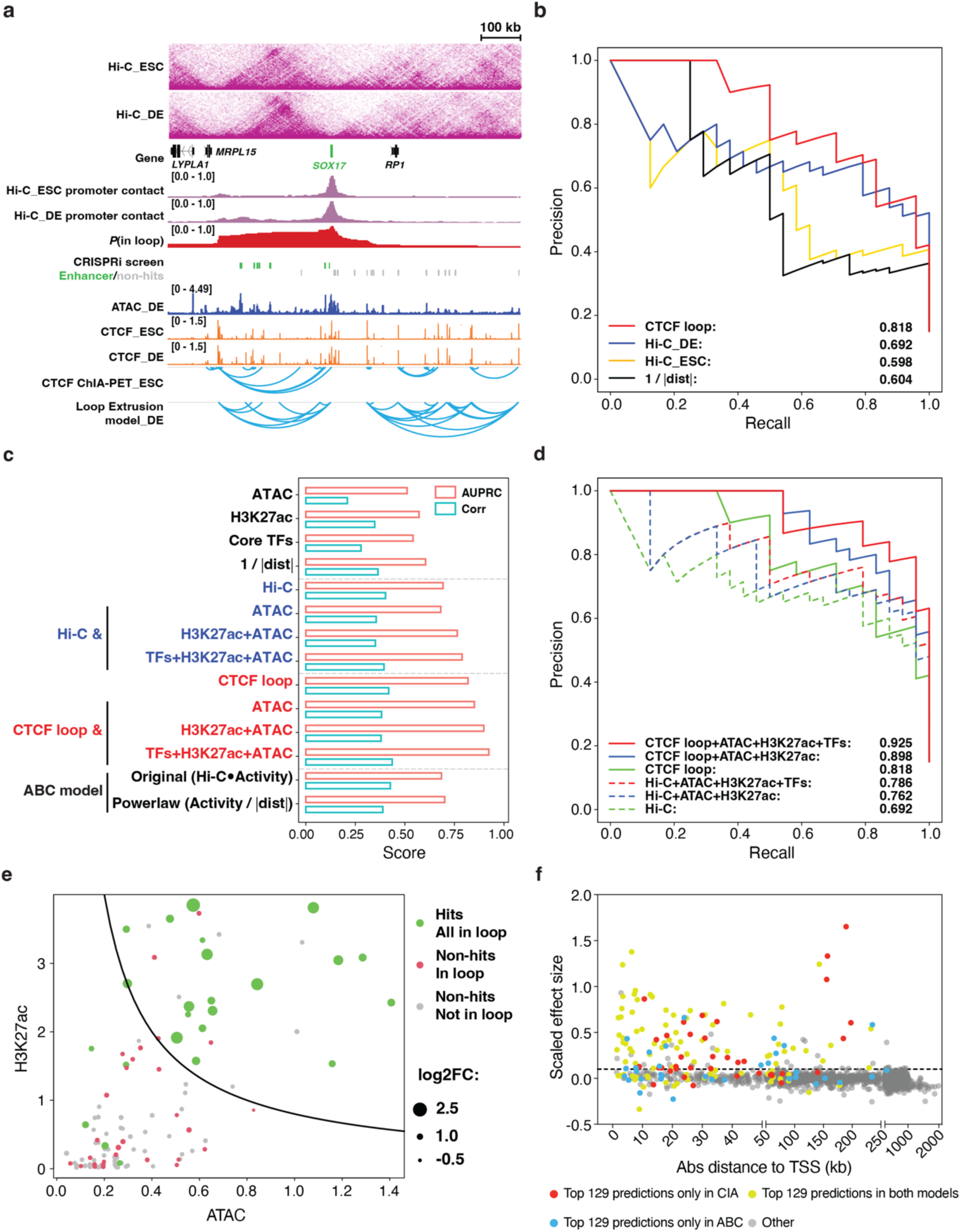
The CIA model provides improved enhancer prediction. **a**, Hi-C-based and CTCF loop-based chromatin conformation analysis at the *SOX17* locus. **b**, A precision-recall plot comparing the performance for prediction of enhancer hits from the screen using *P*(in loop, red), DE Hi-C (blue), ESC Hi-C (yellow) and enhancer-promoter distance (black). **c**, Bar plots comparing the AUPRC and correlation scores between single chromatin feature-based, Hi-C-based, or CTCF loop-based enhancer prediction model with the ABC model. **d**, A precision-recall plot comparing the performance for prediction of enhancer hits from the screen using CTCF loop (solid line) and Hi-C (dash line) with or without additional chromatin feature combination. **e**, A scatter plot showing the combinatory criteria of *P*(in loop), H3K27ac and ATAC can clearly separate the hits (green) and non-hits (red and gray). *P*(in loop) > 0.5 is used to justify the enhancers and targeting promoters are in the same CTCF loop (green and red). The solid line represents the threshold value (criteria) of *Activity* = √*ATAC* ∗ *H*3*K*27*ac*. The size of each dot represents the log2FC of each enhancer from the screen. **f**, A scatter plot showing the comparison between the CIA and ABC model with all 3 datasets (ESC-DE, K562 Reilly^9^ and K562 Nasser^41^). The numbers of selected top predictions are based on the hits identified from each data set, including top 24 predictions from ESC-DE, top 36 predictions from K562 Reilly and top 69 predictions from K562 Nasser. Yellow dots represent top predictions in both models. Red dots represent top predictions only in the CIA model. Cyan dots represent top predictions only in the ABC model. Grey dots represent regions that do not belong to top predictions. The effect sizes from each data set are scaled to reach to the same threshold (∼0.1 dashed line).

## Acknowledgement

We acknowledge the assistance from the following Memorial Sloan Kettering Cancer Center (MSKCC) Cores: Antibody & Bioresource, Molecular Cytogenetics, Gene Editing & Screening, and Integrated Genomics Operation. We thank R. Garippa, H. Liu and S. Mehta for assistance with CRISPR library generation and HiSeq for quantifying gRNA abundance after CRISPRi screen; Y. Furuta, J. Liu, Y. Lan and C. Schwarz for assisting with additional experiments not included in the manuscript; and C.S. Leslie, L. Studer, A. Ventura, A.-K. Hadjantonakis, T. Evans and members of D.H.’s laboratory for insightful advice.

## Author contributions

R.L., D.H. and M.B. devised experiments and interpreted results. R.L. performed most experiments and analyzed the results. M.B. developed the mathematical models and performed computational data analysis, with contributions from J.O., W.X. and D.S.. J.Y., B.R., D.Y. and Q.L. assisted with ChIP-seq. S.S. supervised and J.Y. and R.C. assisted with ChIP-MS. T.V. and D.H. supervised and R.G. and T.C. assisted with validation. E.A. and D.H. supervised and J.P., D.M. and W.W. assisted with Hi-C and subsequent data analysis. H.C. performed gene expression correlation analysis. R.L., D.H. and M.B. wrote the manuscript; all other authors provided editorial advice.

## Data availability

The parental HUES8 hESC line was obtained from Harvard University under a material transfer agreement. Sequencing data are available at GEO, accession no. GSE213394 (new data from this study) and GSE114102 (published DE-72h H3K4me1 ChIP-seq data). Mass spectrometry raw data are available at the free online repository Chorus (https://chorusproject.org/) under project no. 1787.

## Code availability

Code used in this study will be available upon request.

## Extended Data Figures

**Extended Data Figure 1.**
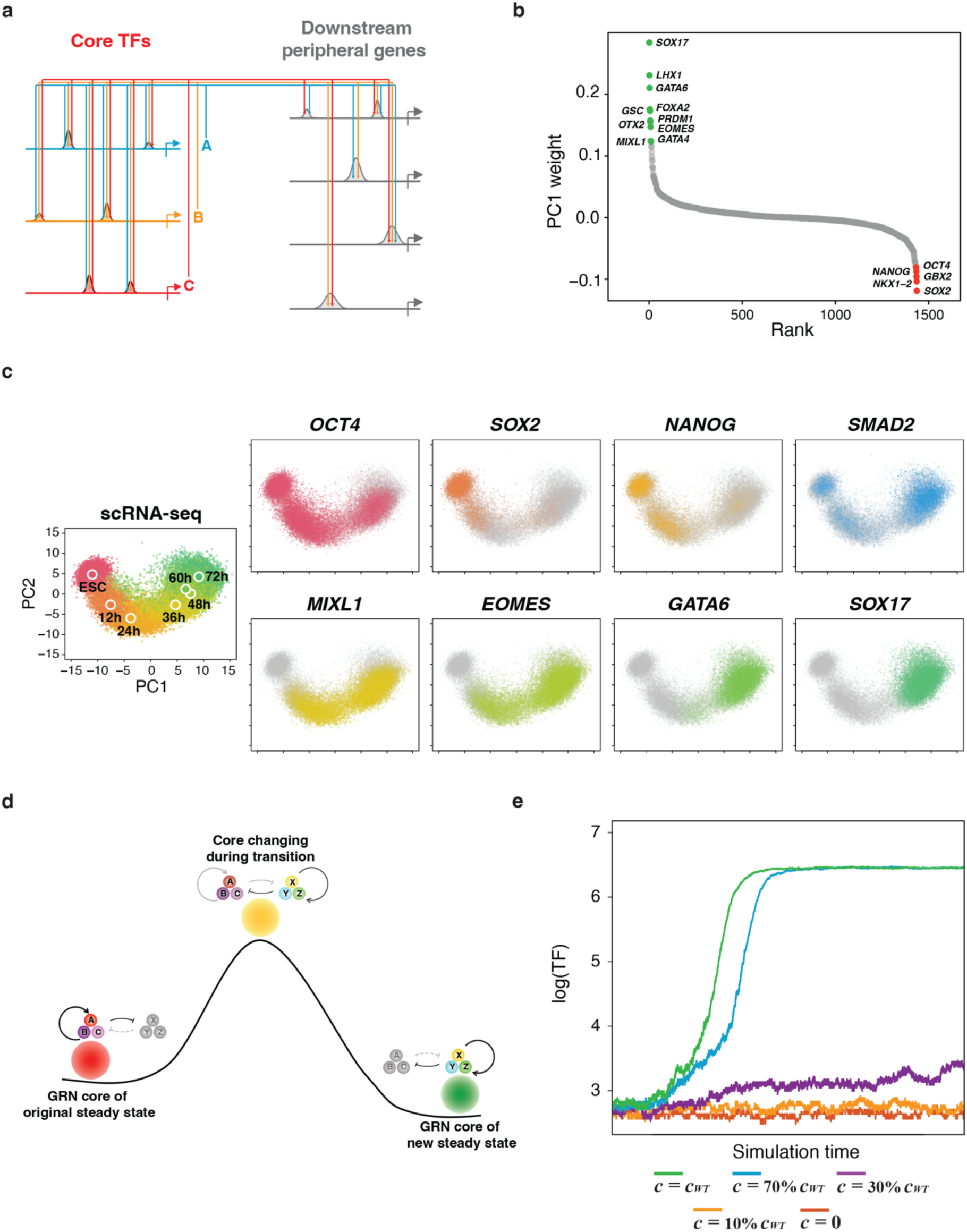
Supporting data for the Dynamic GRN model. **a**, The detailed schematic of core circuit in the GRN. The core TFs cooperatively auto-regulate each other by binding to core enhancers and co-regulate downstream peripheral genes by binding to peripheral enhancers. **b**, The ranking plot of PC1 weight of all TFs in PCA analysis from scRNA-seq data during hESC-DE transition. **c**, The PCA plots showing selective TFs from the PCA component 1 (Extended Data Fig.1b) of scRNA-seq sampled every 12 hours during hESC-DE transition. **d**, The schematic of core circuit establishment during cell state transition, similar to *Moris et al* ^68^. The transition of a cell from one steady state to another is accompanied by the deconstruction of the original core circuit (A, B, C) and the establishment of core circuit of the new state (X, Y, Z). **e**, Stochastic Gillespie simulations of the dynamic GRN network model. The green, cyan, purple, yellow and orange lines represent 100%, 70%, 30%, 10% and 0% of original total enhancer strength respectively.

**Extended Data Figure 2.**
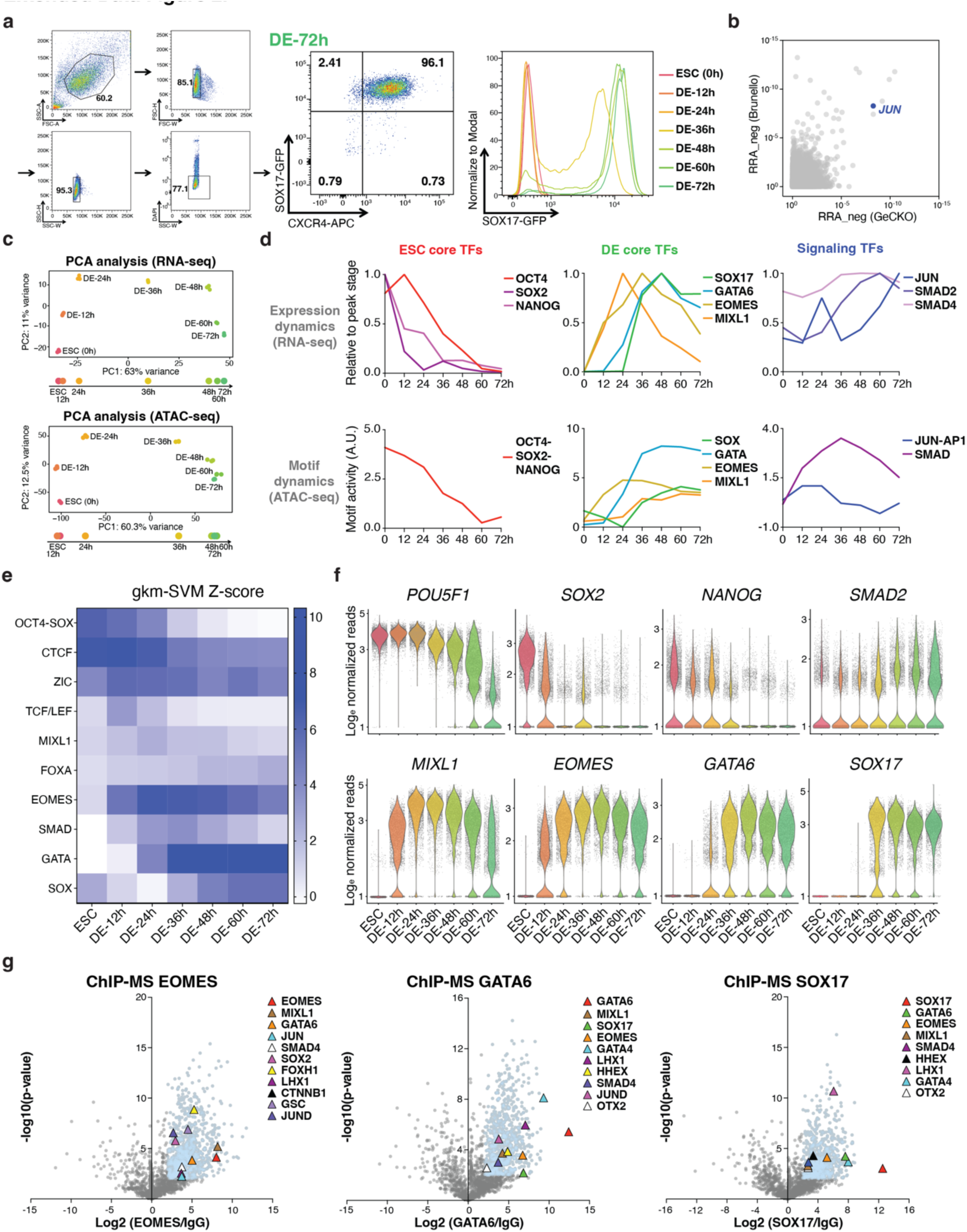
Core TFs identification and characterization during hESC-DE transition. **a**, Flow cytometry analysis showing the gating strategy (left), differentiation efficiency at DE-72h measured by DE markers SOX17 and CXCR4 (middle) or transition efficiency every 12h measured by SOX17 (right). **b**, MAGeCK RRA scores for negative hits in two genome-scale DE screens from Li et al ^25^. JUN is the only identified TF among the negative hits. **c**, PCA analysis of bulk RNA-seq (top) and ATAC-seq (bottom) during hESC-DE transition. Data points are projected to PC1 (cell state) to determine the time window of cell state transition. **d**, The expression dynamics (top) and motif dynamics (predicted by gkm-SVM trained on the ATAC-seq data, bottom) of core TFs during hESC-DE transition. **e**, Motif Z-score of ATAC-seq by gkm-SVM at each time point during hESC-DE transition. **f**, Feature violin plots from scRNA-seq data showing core TFs expression changing during hESC-DE transition at single cell resolution. **g**, Volcano plots showing protein-protein interactions identified by ChIP-MS using EOMES as the bait at DE-24h, GATA6 and SOX17 as baits at DE-48h. Blue dots represent the significantly enriched proteins with log2FC > 2 and -log10(p-value) > 2. Selective TFs enriched in ESC and endoderm GO terms are labeled by triangles.

**Extended Data Figure 3.**
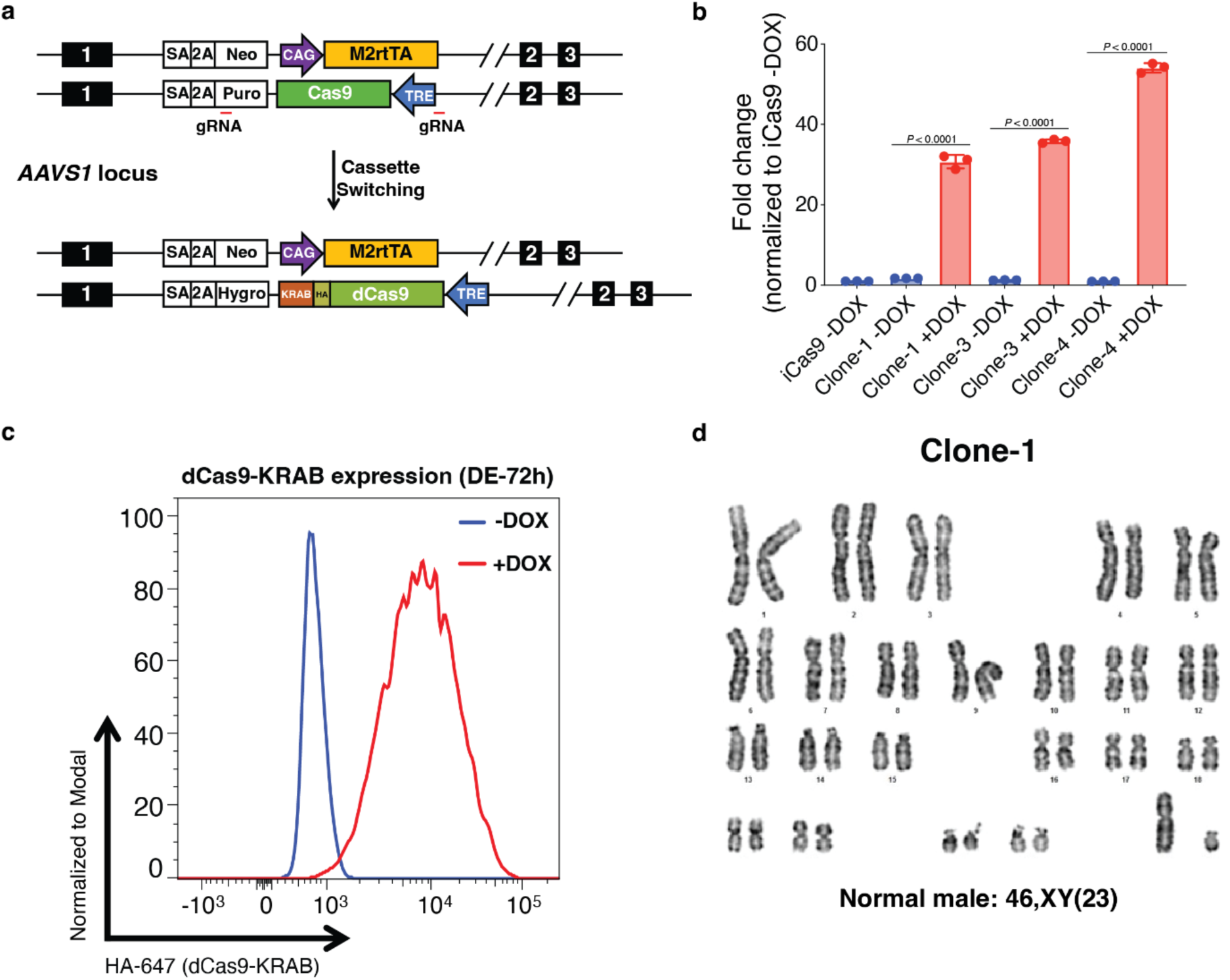
idCas9-KRAB *SOX17^GFP/+^* hESC line generation using cassette switching. **a**, The schematics of idCas9-KRAB *SOX17^GFP/+^* hESC line generation using cassette switching. gRNAs targeting the puromycin selection cassette and the 5’ sequence outside TRE are designed for inducing double-strand break for homology repair. **b**, RT-qPCR results showing the inducible expression of dCas9-KRAB with doxycycline treatment. n=3 independent experiments. Error bars indicate mean ± SD. Statistical analysis was performed by two-tailed unpaired student t-test. **c**, Flow cytometry results showing the inducible expression of dCas9-KRAB with doxycycline treatment. **d**, Karyotyping results of the idCas9-KRAB *SOX17^GFP/+^* hESC line.

**Extended Data Figure 4.**
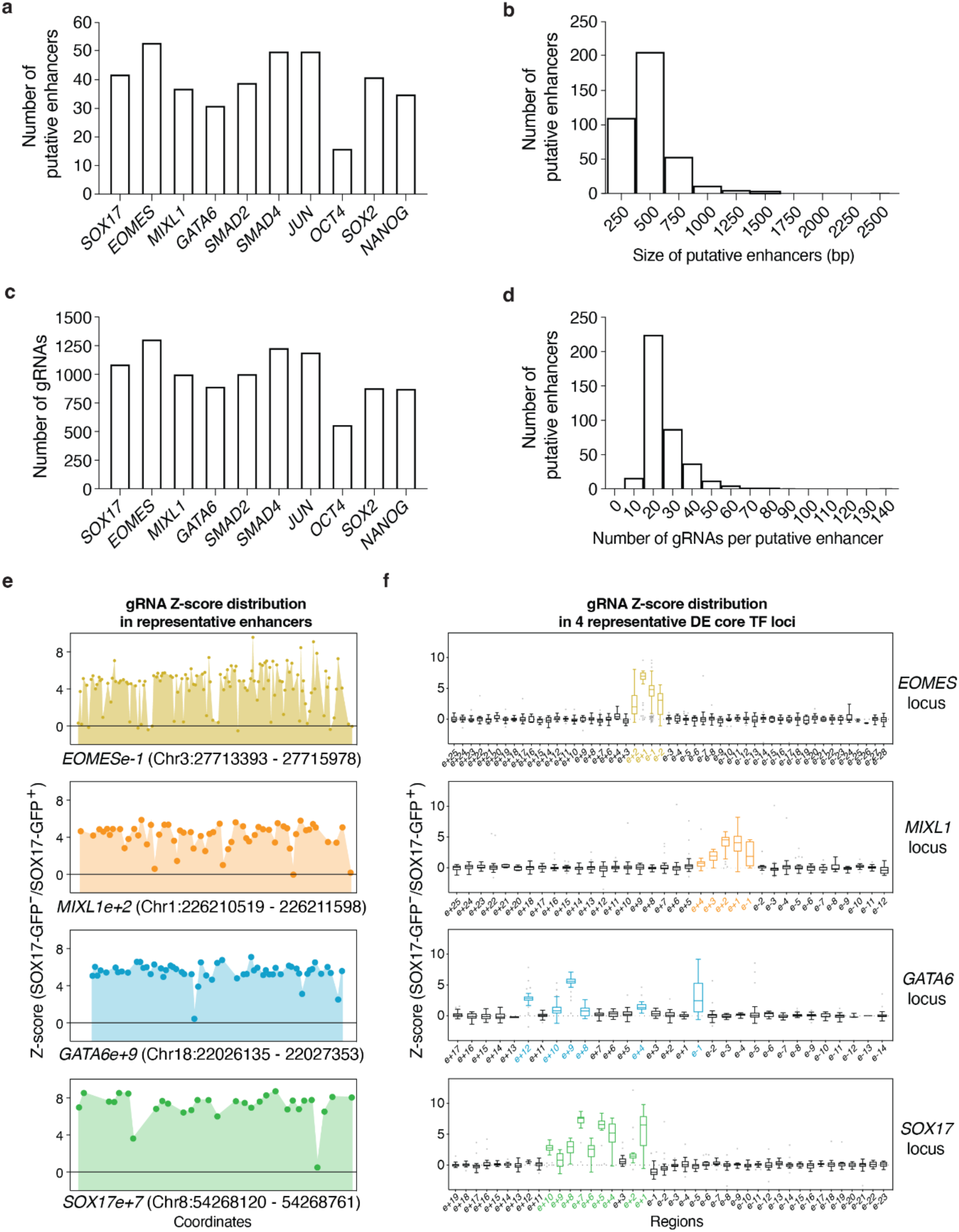
Supporting data for the screen design and gRNA enrichment analysis. **a** to **d**, Statistics of putative enhancers selection and gRNAs design. The number of putative enhancers selected for each core TFs (**a**), the size of putative enhancers (**b**), the total number of gRNAs targeted on putative enhancers of each core TF (**c**), the number of gRNAs targeted on each putative enhancer (**d**). **e**, The gRNA Z-score distribution at representative enhancers showing gRNAs targeting the same enhancer have similar perturbation effect. **f**, Box plots showing the gRNA Z-score distribution in all putative enhancers of *EOMES*, *MIXL1, GATA6* and *SOX17* loci. Center lines indicate median. Boxes limit upper and lower quartiles.

**Extended Data Figure 5.**
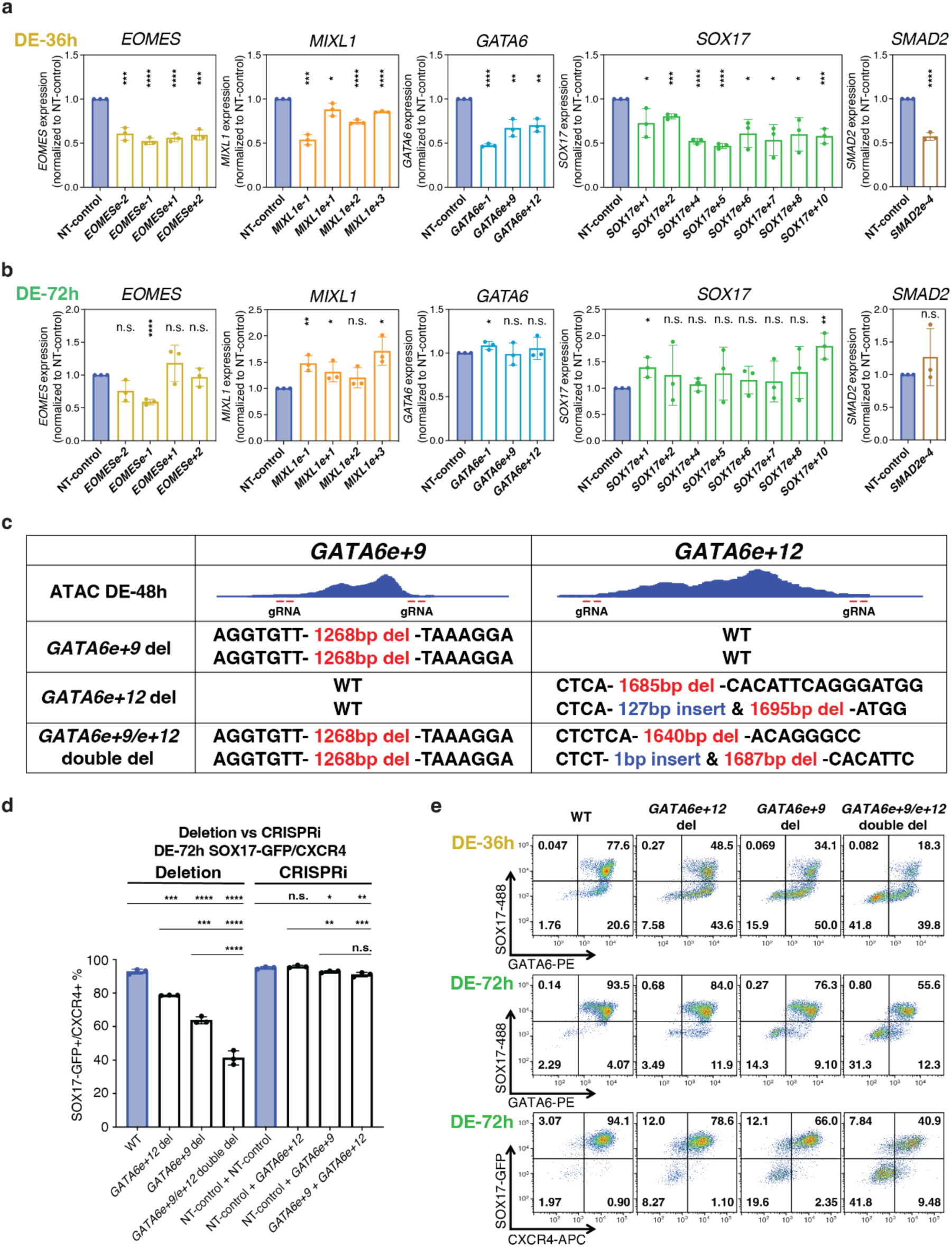
Supporting data for validation of core enhancers. **a**-**b**, RT-qPCR showing the expression of the cognate genes decreases by enhancer perturbations at DE-36h (**a**) but mostly restored at DE-72h (**b**). n=3 independent experiments. Error bars indicate mean ± SD. Statistical analysis was performed by two-tailed unpaired student t-test. Significance is indicated as: *P < 0.05, **P < 0.01, ***P < 0.001 and ****P < 0.0001. n.s.: not significant. **c**, Illustration of the enhancer deletion experiments that resulted in the *GATA6e+9* deletion (del), *GATA6e+12* del and *GATA6e+9/e+12* double del hESC lines. **d**, Statistics of SOX17-GFP/CXCR4 double positive cells at DE-72h in WT, *GATA6e+9* del, *GATA6e+12* del, *GATA6e+9/e+12* double del cells, as well as cells with non-targeting control, *GATA6e+9* perturbation, *GATA6e+12* perturbation and *GATA6e+9/GATA6e+12* dual-perturbation. n=3 biological replicates. Error bars indicate mean ± SD. Statistical analysis was performed by two-tailed unpaired multiple comparison test with Dunnett correction. Significance is indicated as: *P < 0.05, **P < 0.01, ***P < 0.001 and ****P < 0.0001. n.s. Not significant. **e**, Flow plots showing SOX17/GATA6 expression at DE-36h, DE-72h and SOX17-GFP/CXCR4 expression at DE-72h of WT, *GATA6e+9* del, *GATA6e+12* del, *GATA6e+9/e+12* double del.

**Extended Data Figure 6.**
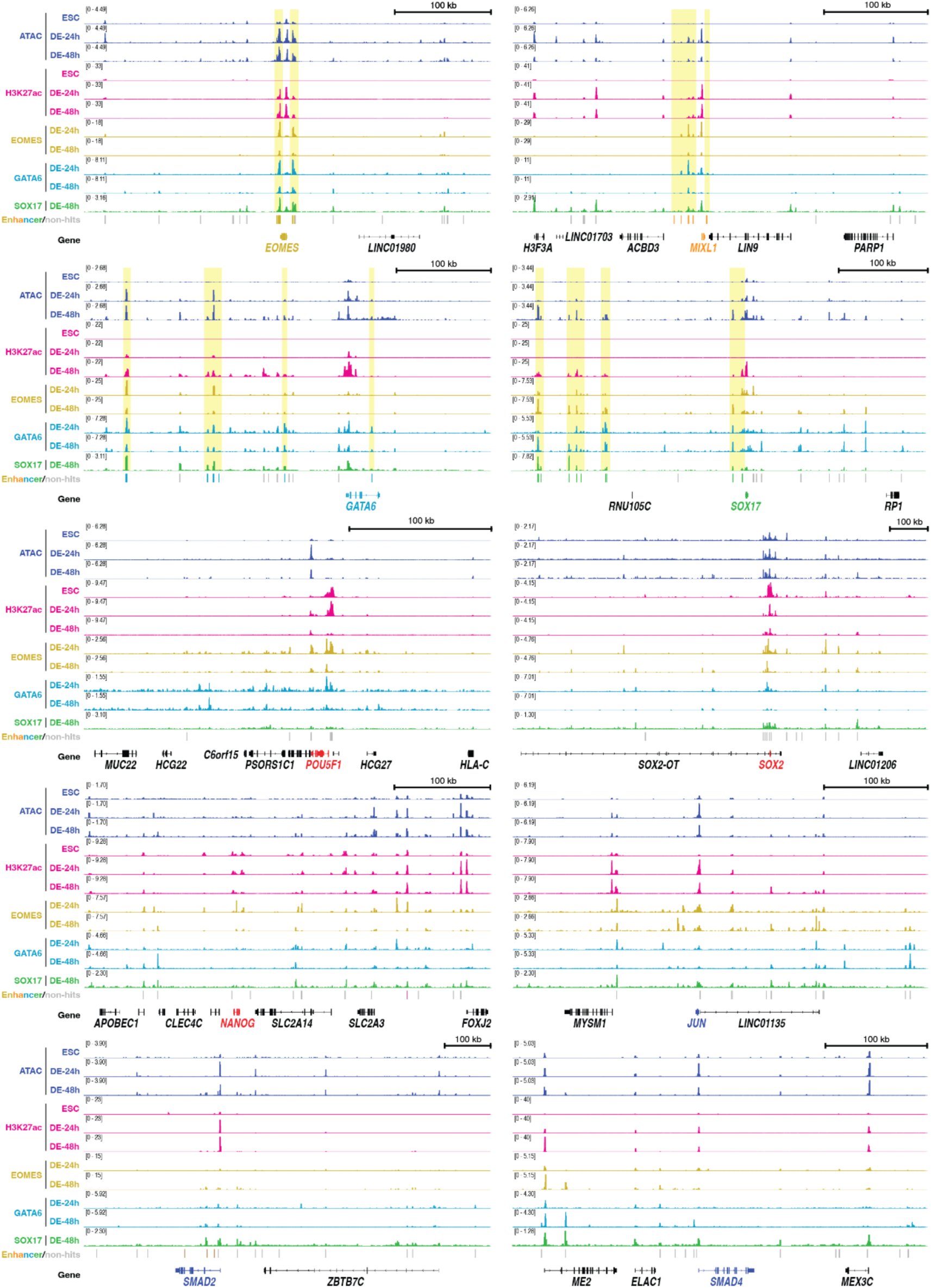
Epigenetic features of core enhancers. Relevant ATAC-seq and ChIP-seq tracks of all 10 selected loci. Yellow boxes highlight the DE core TFs (EOMES, GATA6 and SOX17) bind to DE core enhancers. Genomic coordinates from GRCh38 (human hg38) for each gene are labeled. kb, kilobase.

**Extended Data Figure 7.**
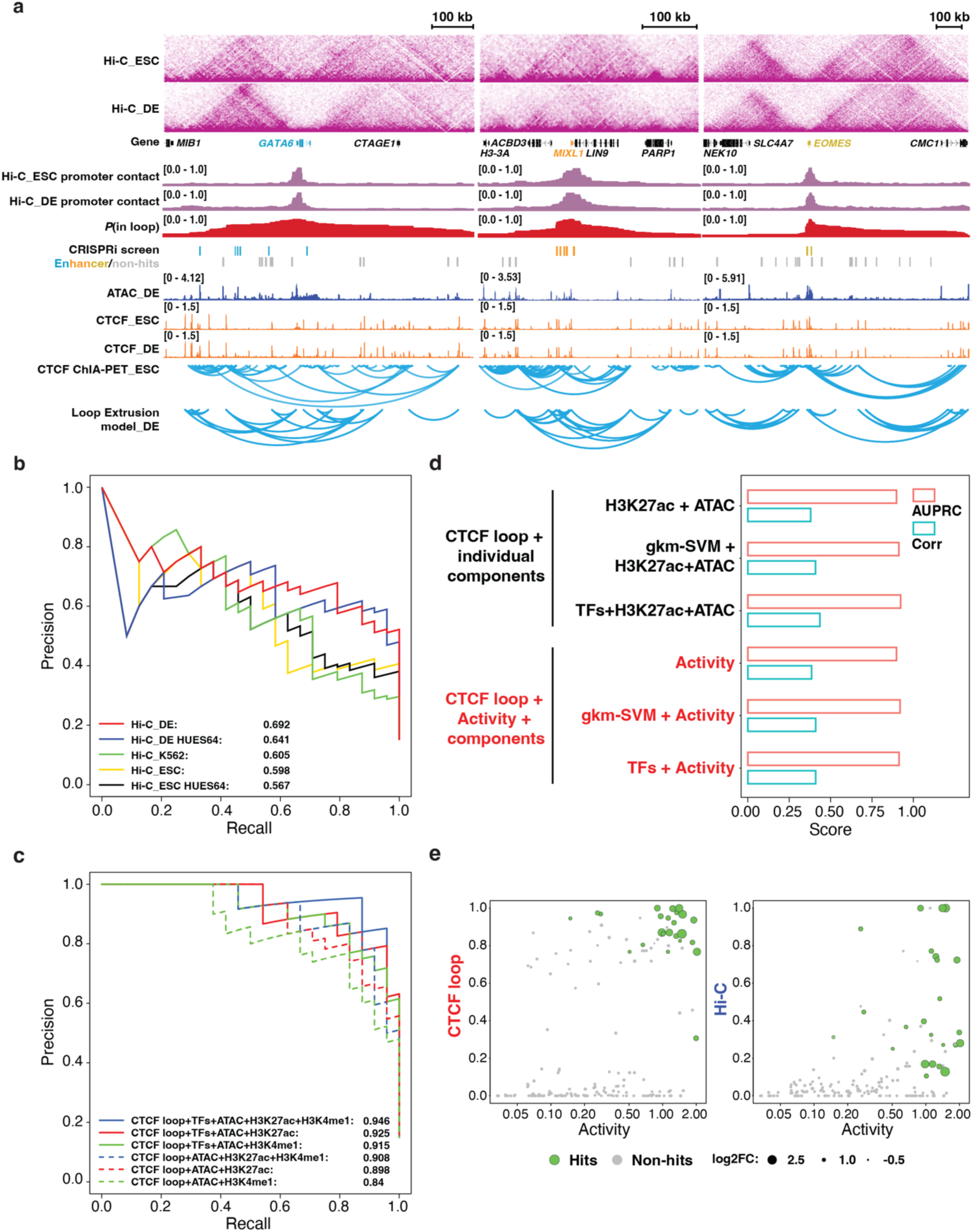
Supporting data for enhancer prediction using CIA model. **a**, Hi-C-based and CTCF loop-based chromatin conformation analysis at the *GATA6*, *MIXL1* and *EOMES* loci. **b**, Precision-recall plot comparing the performance for prediction of enhancer hits from the screen using different Hi-C datasets. **c**, Precision-recall plot comparing the performance for prediction of enhancer hits from the screen using CIA model with additional H3K4me1 chromatin feature. **d**, Bar plot comparing the AUPRC and correlation scores between logistic regression of chromatin feature combination and 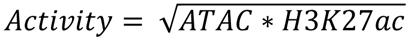 in CIA model. **e**, A scatter plot showing the *P*(in loop) can classify hits (green) and non-hits (grey) more clearly than Hi-C-based enhancer-promoter contact frequency. The size of each dot represents the log2FC of each enhancer from the screen.

**Extended Data Figure 8.**
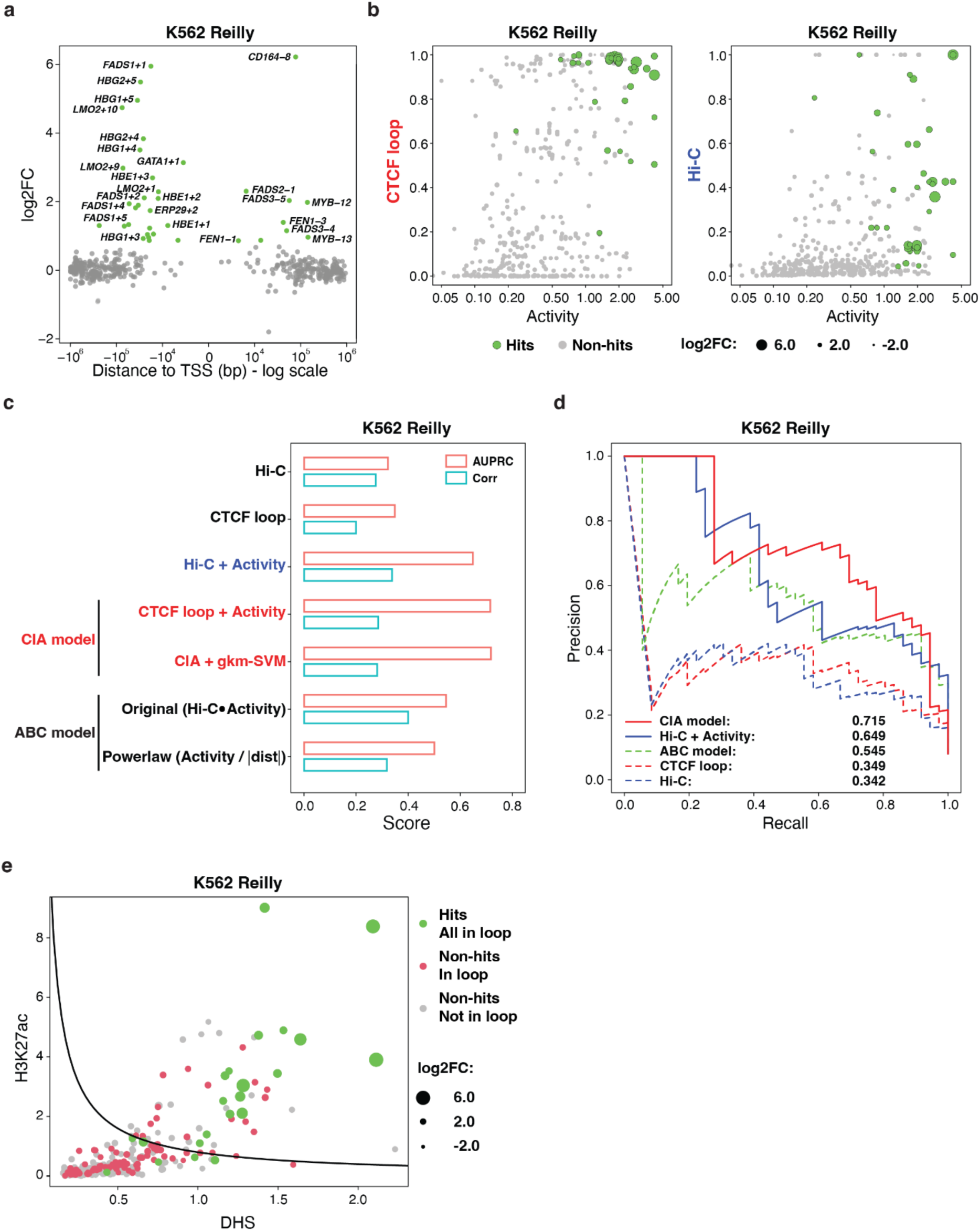
The CIA model predicts active enhancers in different scenarios (K562 Reilly). **a**, gRNA enrichment analysis identified 36 hits from the HCR-FF screen in K562 cells from *Reilly et al* ^9^ **b**, The scatter plot showing the *P*(in loop) can classify hits (green) and non-hits (grey) in K562 HCR-FF screen more clearly than Hi-C-based enhancer-promoter contact frequency. The size of each dot represents the log2FC of each enhancer from the screen. **c**, Bar plot comparing the AUPRC and correlation scores between Hi-C-based enhancer prediction with CIA model and ABC model using K562 HCR-FF screen results. **d**, Precision-recall plot comparing the performance for prediction of enhancer hits from the K562 HCR-FF screen using CTCF loop-based model and Hi-C-based model. **e**, The scatter plot showing the combinatory criteria of *P*(in loop), H3K27ac and ATAC can clearly separate the hits (green) and non-hits (red and gray) from the K562 HCR-FF screen. *P*(in loop) > 0.5 is used to highlight enhancers and targeting promoters in the same CTCF loop (green and red). The solid line represents the same threshold criterion of 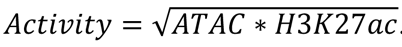 in Fig.4e. The size of each dot represents the log2FC of each enhancer from the K562 HCR-FF screen.

**Extended Data Figure 9.**
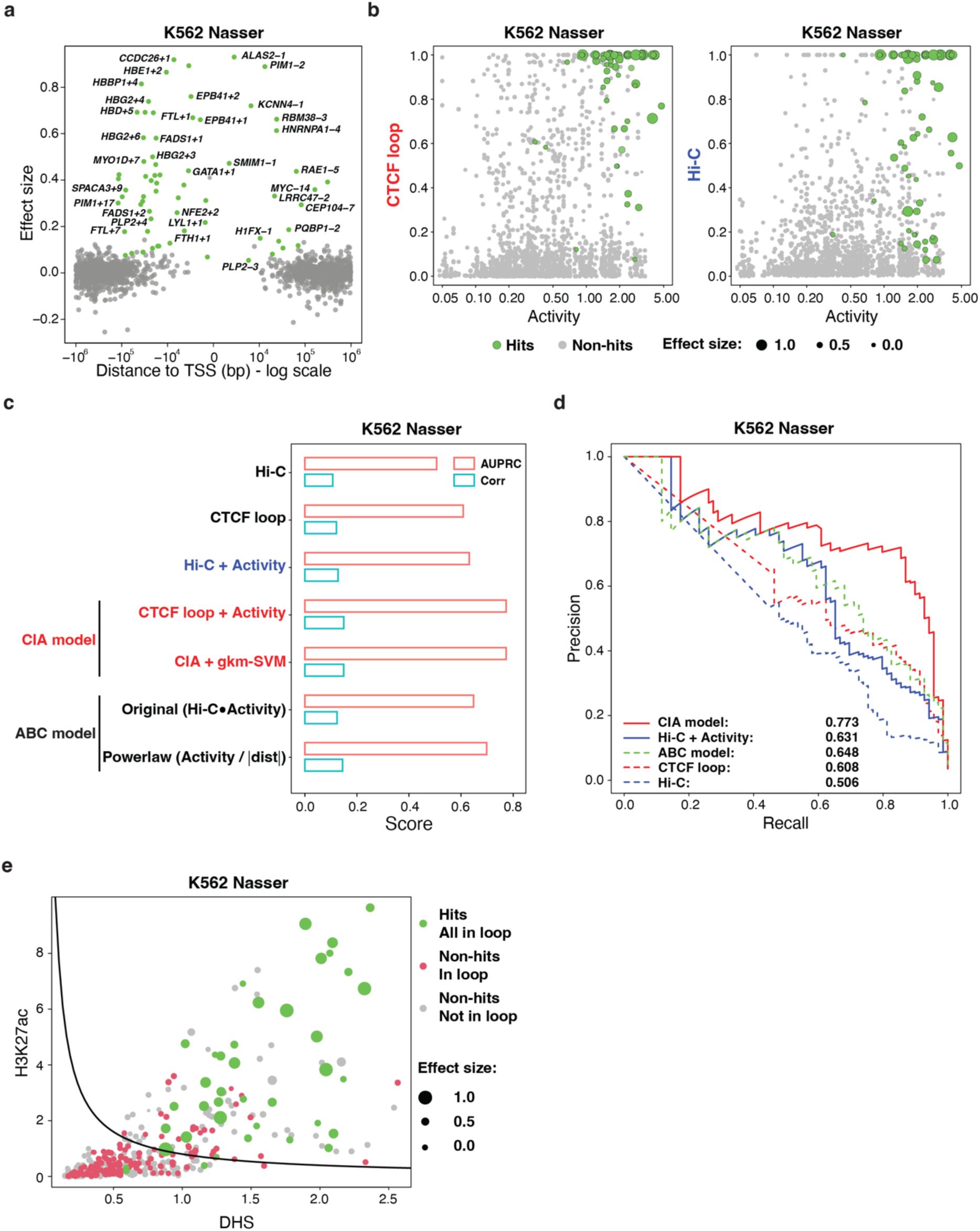
The CIA model predicts active enhancers in different scenarios (K562 Nasser). **a**, 69 identified hits in the K562 cells from *Nasser et al* are plotted^41^. **b**, The scatter plot showing the *P*(in loop) can classify hits (green) and non-hits (grey) in K562 Nasser more clearly than Hi-C-based enhancer-promoter contact frequency. The size of each dot represents the effect size of each enhancer from *Nasser et al*. **c**, Bar plot comparing the AUPRC and correlation scores between Hi-C-based enhancer prediction with CIA model and ABC model using K562 Nasser results. **d**, Precision-recall plot comparing the performance for prediction of enhancer hits from the K562 Nasser using CTCF loop-based model and Hi-C-based model. **e**, The scatter plot showing the combinatory criteria of *P*(in loop), H3K27ac and ATAC can clearly separate the hits (green) and non-hits (red and gray) from the K562 Nasser. *P*(in loop) > 0.5 is used to highlight enhancers and targeting promoters in the same CTCF loop (green and red). The solid line represents the same threshold criterion of 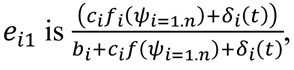 in Fig.4e. The size of each dot represents the effect size of each enhancer from the K562 Nasser.

## Extended Data

**Extended Data Table 1.** EOMES, GATA6 and SOX17 ChIP-MS data.

**Extended Data Table 2.** Core enhancer perturbation screen gRNA library sequences and gRNA enrichment analysis.

**Extended Data Table 3.** Core enhancer perturbation screen gRNA enrichment analysis for regions.

**Extended Data Table 4.** Data of core enhancer screening for regions surrounding core DE TFs.

**Extended Data Table 5.** Data of K562 Reilly enhancer screen.

**Extended Data Table 6.** Data of K562 Nasser enhancer screen.

**Extended Data Table 7.** gRNA for cell line generation.

**Extended Data Table 8.** Primer sequences for cell line genotyping, RT-qPCR and gRNA enrichment analysis.

**Extended Data Table 9.** Antibody for flow cytometry, ChIP-seq and ChIP-MS.

**Extended Data Table 10.** gRNA sequences for hit validation.

